# Unexpected roles for AMPK in the suppression of autophagy and the reactivation of mTORC1 signaling during prolonged amino acid deprivation

**DOI:** 10.1101/2023.12.20.572593

**Authors:** Dubek Kazyken, Sydney G. Dame, Claudia Wang, Maxwell Wadley, Diane C. Fingar

**Author notes:** Co-corresponding authors: Diane C. Fingar, Ph.D. and Dubek Kazyken, Dept. of Cell and Developmental Biology, University of Michigan Medical School, BSRB #3039, 109 Zina Pitcher Place, Ann Arbor, MI 48109-2200. (dcf office); (lab).

## Abstract

AMPK promotes catabolic and suppresses anabolic cell metabolism to promote cell survival during energetic stress, in part by inhibiting mTORC1, an anabolic kinase requiring sufficient levels of amino acids. We found that cells lacking AMPK displayed increased apoptotic cell death during nutrient stress caused by prolonged amino acid deprivation. We presumed that impaired autophagy explained this phenotype, as a prevailing view posits that AMPK initiates autophagy (often a pro-survival response) through phosphorylation of ULK1. Unexpectedly, however, autophagy remained unimpaired in cells lacking AMPK, as monitored by several autophagic readouts in several cell lines. More surprisingly, the absence of AMPK increased ULK1 signaling and LC3b lipidation during amino acid deprivation while AMPK-mediated phosphorylation of ULK1 S555 (a site proposed to initiate autophagy) decreased upon amino acid withdrawal or pharmacological mTORC1 inhibition. In addition, activation of AMPK with compound 991, glucose deprivation, or AICAR blunted autophagy induced by amino acid withdrawal. These results demonstrate that AMPK activation and glucose deprivation suppress autophagy. As AMPK controlled autophagy in an unexpected direction, we examined how AMPK controls mTORC1 signaling. Paradoxically, we observed impaired reactivation of mTORC1 in cells lacking AMPK upon prolonged amino acid deprivation. Together these results oppose established views that AMPK promotes autophagy and inhibits mTORC1 universally. Moreover, they reveal unexpected roles for AMPK in the suppression of autophagy and the support of mTORC1 signaling in the context of prolonged amino acid deprivation. These findings prompt a reevaluation of how AMPK and its control of autophagy and mTORC1 impact health and disease.

## Introduction

The conserved processes of macroautophagy and selective autophagy (e.g. mitophagy) play important roles in protein quality control, cell homeostasis, and cell survival [1–3]. During macroautophagy (hereafter, autophagy), bulk cytoplasm containing proteins and organelles become engulfed in double-membrane structures called autophagosomes that fuse with lysosomes to form autolysosomes, which degrade and thereby recycle sequestered cargo. A complex set of autophagy (ATG) proteins orchestrates this catabolic response to maintain proteostasis in healthy cells, provide metabolic building blocks during amino acid deprivation, and remove damaged proteins and organelles during stress or disease [1–3]. As such, defective autophagy contributes to various diseases including the neurodegenerative disorders Alzheimer Disease (AD) and Amyotrophic Lateral Sclerosis (ALS), which arise in part due to the accumulation of pathological protein aggregates [4]. Elevated autophagy has also been linked to cancer, as it provides metabolites and energy that fuel cell metabolism, growth, and proliferation [4].

mTOR complex 1 (mTORC1) promotes anabolic cellular processes (e.g., ribosomal biogenesis; protein, lipid, and nucleotide synthesis) in the presence of sufficient levels of nutrients (amino acids; glucose), growth factors, and hormones [5–8]. Well-characterized substrates of mTORC1 include S6K1 and 4EBP1, which control cap-dependent translation [5–8]. mTORC1 also suppresses the initiation of autophagy in nutrient- and growth factor-rich environments by phosphorylating and inhibiting the kinase ULK1 [9,10] (ATG1 in yeast). Amino acid deprivation inactivates mTORC1 rapidly (30-60 minutes), which initiates autophagy by relieving the suppressive effect of mTORC1 on ULK1 [11–13]. At later time points of amino acid withdrawal, mTORC1 re-activates partially (∼2-4 hr, depending on cell type) due to the induction of autophagy and the subsequent liberation of internal amino acids by autophagic proteolysis [11,14–16]. This reactivation of mTORC1 limits the extent of autophagy (which can induce cell death) and enables the regeneration of lysosomes, a process known as ALR (autophagic lysosome regeneration) [14,17].

ULK1 nucleates a multi-protein complex containing ATG13 and FIP200 (FAK family-interacting protein of 200 kDa) that initiates autophagy [1–3,18]. The activated ULK1 complex next activates a complex containing Vps34 (a class III phosphatidylinositol-3-kinase) through phosphorylation of its partners ATG14 and Beclin-1. The Vps34 complex generates PtdIns3P (phosphatidylinositol-3-phosphate) on ATG9-containing membrane vesicles. These vesicles nucleate the formation of double membrane phagophores, which expand around cargo to form mature autophagosomes. PtdIns3P on growing phagophores recruits WIPI2 proteins and ATG16L1-ATG5-ATG12 complexes, which covalently attach LC3 proteins (ATG8 in yeast) to the lipid phosphatidylethanolamine (PE) on growing phagophore membranes, a process known as LC3 lipidation. Indeed, the most widely employed assay for monitoring autophagy measures the conversion of LC3b (LC3b-I) to a lipidated form known as LC3b-II that migrates faster on SDS-PAGE I [19]. This assay can be misinterpreted, however, if not performed rigorously by utilizing autophagic flux inhibitors (e.g., BafA1, an H^+^-vATPase inhibitor that maintains an acidic pH within lysosomes; inhibitors of acid-dependent proteases; or NH_4_Cl, an agent that alkalinizes lysosomes) [19,20]. These flux inhibitors blunt late-stage autophagic proteolysis of lipidated LC3b-II and autophagy receptors such as p62/SQSTM1, thus enabling an accurate assessment of LC3b lipidation (i.e., LC3b-II) and p62/SQSTM1 levels in the absence of degradative flux [19].

Induction of energetic stress (i.e., a rise in the AMP or ADP to ATP ratio) caused by prolonged glucose deprivation or mitochondrial poisons inhibits mTORC1 signaling by activating AMPK (AMP-activated protein kinase) (Snf1p in yeast) [11,21–27]. Mild glucose withdrawal can also activate AMPK in an ATP-independent manner [28]. AMPK activation redirects cell metabolism away from ATP-consuming anabolic processes and toward ATP-generating, catabolic processes (e.g., fatty acid oxidation; glucose uptake; lysosomal and mitochondrial biogenesis) that in turn balance energy supply and demand and therefore promote cell survival [11,24,25,27]. AMPK functions as a heterotrimer composed of a catalytic α-subunit (α1−2) and regulatory β- and γ-subunits (β1-2; γ1-3) [24,25,27]. Due to its heterotrimeric assembly and multiple subunits, AMPK can assemble into 12 distinct isoforms [24,25,27], although most subunits display tissue-specific expression and thus most cells express only a subset of these AMPKαβγ isoforms [25]. During energetic stress, AMP (or ADP) binds to the γ-subunit, leading to a conformational change that increases the phosphorylation of the catalytic AMPK α-subunit on its essential activation loop site (T172) by the tumor suppressor LKB1 or the Ca^2+^-requiring kinase CaMKKβ [24,25,27].

A widely accepted model posits that activation of AMPK by glucose deprivation induces autophagy and mitophagy through multi-site phosphorylation of ULK1 and Beclin-1 [2,3,9,27,29–33]. In addition, AMPK-mediated phosphorylation of Tsc2 [21] and Raptor (an essential mTORC1 partner protein) inhibits signaling mTORC1 in response to energetic stress [21,22], which theoretically upregulates autophagy. In this prevailing model, mTORC1 phosphorylates ULK1 S757 (human S758) in complete media (i.e., contains glucose and amino acids), which destabilizes a complex between AMPK and ULK1, thereby reducing AMPK-mediated phosphorylation and activation of the ULK1 complex [29]. Conversely, the model proposes that activation of AMPK upon glucose withdrawal reduces mTORC1-mediated phosphorylation of ULK1 S757, which stabilizes the AMPK-ULK1 complex, thus enabling AMPK to phosphorylate ULK1 on several sites, resulting in the initiation of autophagy [29,30]. These findings have led to the widely accepted idea that AMPK promotes autophagy [2,3,27,34]. It is important to note, however, that several reports have questioned aspects of this model. For example, several studies found that glucose withdrawal failed to induce autophagy in mammalian cells and yeast [35–37] while glucose withdrawal suppressed autophagy induced by amino acid withdrawal [37]. In addition, the withdrawal of amino acids or pharmacological inhibition of mTORC1, conditions that induce autophagy, destabilized the AMPK-ULK1 complex and reduced the phosphorylation of ULK1 S555 [10,37–39], a site phosphorylated by AMPK and proposed to induce autophagy [30,32]. Thus, conundrums exist regarding how mTORC1 activity controls the stability of the AMPK-ULK1 interaction and the role of glucose deprivation and AMPK activation in the control of autophagy. Due to the interest in AMPK as a therapeutic drug target for several chronic diseases (e.g., type II diabetes) and its dual role as a tumor suppressor or tumor promoter depending on cellular context [25,40–45], it is important to clarify the role of AMPK in autophagy.

We initiated this study after observing increased apoptotic cell death with intact autophagy in cells lacking AMPK relative to control cells during prolonged amino acid deprivation. At the outset, we presumed that impaired autophagy explained the increased cell death in cells lacking AMPK (as autophagy often promotes cell survival), but we found unexpectedly that autophagy induced by amino acid withdrawal or mTORC1 inhibition remained unimpaired in cells lacking AMPK. More surprisingly, the absence of AMPK increased while the activation of AMPK suppressed autophagy induced by amino acid withdrawal. Finding that AMPK suppresses autophagy in the context of amino acid deprivation challenges the prevailing view that AMPK promotes autophagy universally. Due to the unexpected effect of AMPK loss on autophagy, we were curious as to how AMPK controls mTORC1 signaling in the context of prolonged amino acid deprivation. Surprisingly, we observed that the reactivation of mTORC1 signaling was impaired in cells lacking AMPK in response to the induction of autophagy and the liberation of internal amino acids by autophagic proteolysis. These results oppose the established view that AMPK inhibits mTORC1 universally. As AMPK suppresses autophagy during amino acid deprivation, AMPK therefore supports mTORC1 signaling in this context not by inducing autophagy but by a new mechanism. Together these findings reveal unexpected, non-canonical roles for AMPK in the suppression of autophagy and reactivation of mTORC1 signaling during prolonged amino acid deprivation.

## Results

### Increased apoptotic cell death with unimpaired autophagy in cells lacking AMPK during amino acid deprivation

AMPK promotes cell survival in response to energetic stress [24,25,27,46] in part by directly activating mTOR complex 2 (mTORC2) and its signaling to Akt, a pro-survival kinase [47]. We were thus curious as to whether AMPK also promotes cell survival during nutrient stress caused by amino acid deprivation. We, therefore, examined how amino acid withdrawal for 0.5-16 hr affects apoptosis in MEFs and HeLa cells containing or lacking the AMPK catalytic subunits AMPKα1 and AMPKα2. Markers of apoptosis, i.e., cleavage of PARP or caspase 3, increased modestly in wild type cells at 8-16 hr of amino acid withdrawal but increased more strongly in AMPKα double knockout (DKO) cells (Fig. 1A, 1B). Moreover, AMPKα DKO MEFs displayed reduced cell viability upon amino acid withdrawal for 16 hr, as measured by ReadyProbe staining that distinguishes live from dead cells (Fig. 1C, 1D) [46].

**Figure 1.**
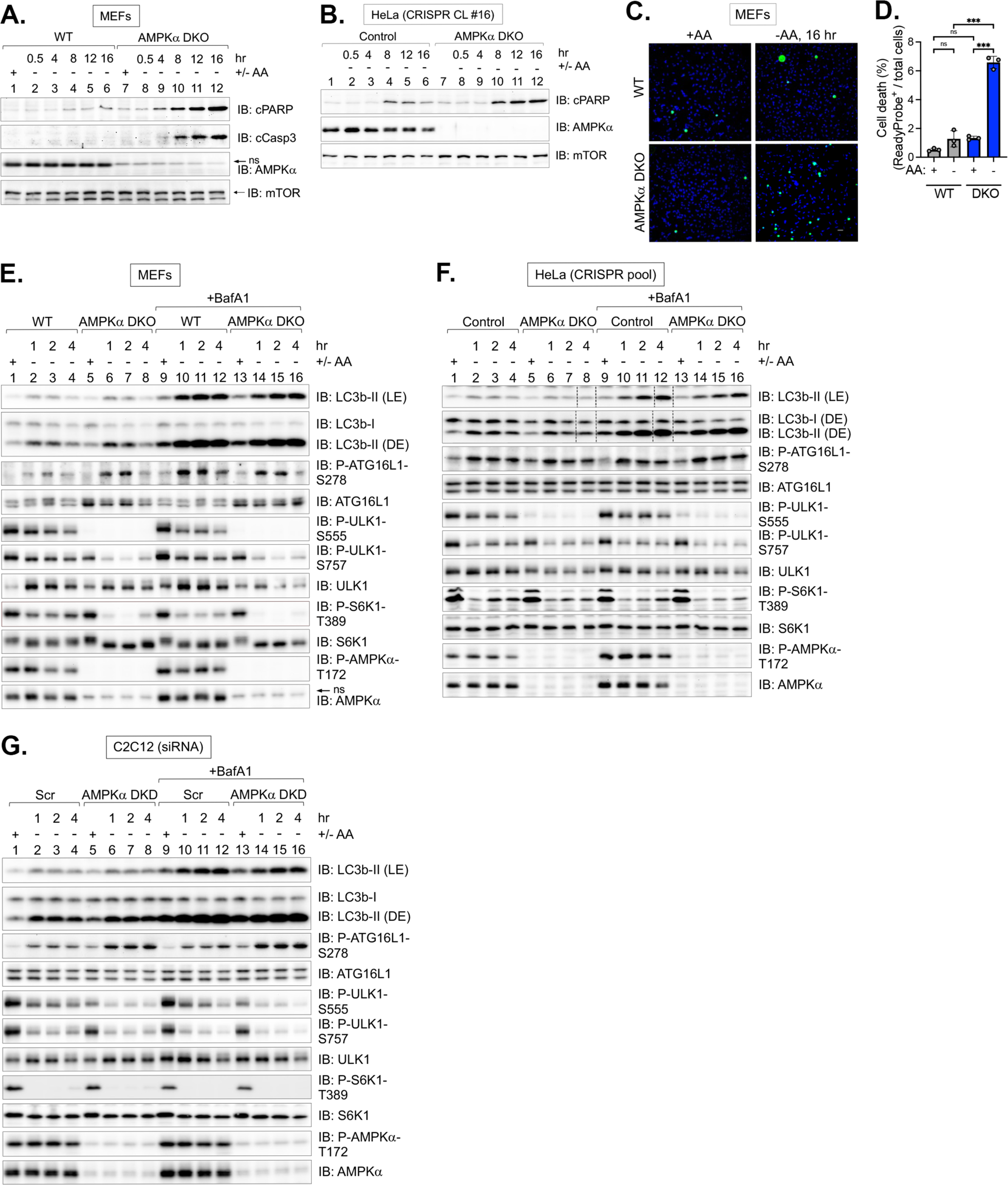
AMPKα1/α2 double knockout (DKO) increases apoptotic cell death and impairs the reactivation of mTORC1 signaling during prolonged amino acid deprivation. **(A)** Immortalized wild type (WT) and AMPKα DKO MEFs isolated from genetically modified mice were fed with complete media (+) for 1 hr or amino acid-free media (−) for 0.5-16 hr. Attached and floating cells were collected, and whole cell lysates were immunoblotted as indicated. ns, non-specific band detected by the AMPKα antibody. **(B)** Control and AMPKα DKO HeLa cells generated by CRISPR/Cas9-mediated genome editing were treated as in (A). **(C)** MEFs were treated as in (A), except ReadyProbe cell viability reagents were added to the culture media. Green fluorescence marks dead cells while blue (Dapi) fluorescence marks total cells. **(D)** Percent (%) cell death is indicated by the ratio of green cells/ total blue cells. n = 100 cells per treatment from 1 representative experiment. Two-way ANOVA was performed followed by Sidak’s multiple comparison post hoc tests; *\*\*\*p* < 0.001; ns, not significant. **(E)** WT and AMPKα DKO MEFs were fed with complete media (+) for 1 hr or amino acid-free media (−) for 1, 2, or 4 hr in the absence or presence of BafA1. Whole cell lysates were immunoblotted as indicated. LE: light exposure; DE: dark exposure. ns, non-specific band detected by the AMPKα antibody in MEFs. **(F)** Control and AMPKα DKO HeLa cells were treated as in (E). Lanes #8 and #12 in the digital LC3b image were re-ordered using Adobe Photoshop, as indicated by thin dotted lines. **(G)** C2C12 cells were transiently transfected with scrambled (Scr) or AMPKα1 and α2 siRNA and treated as in (E).

We presumed that impaired autophagy explained the increased cell death in cells lacking AMPK, as autophagy often promotes cell survival. We, therefore, tested whether the induction of autophagy in response to amino acid deprivation requires AMPK by studying MEFs and other cell lines with double knockout (DKO) of AMPKα1/α2 or C2C12 cells with acute siRNA-mediated double knockdown (DKD) of AMPKα1/α2. To monitor autophagy, we measured the lipidation of LC3b and the phosphorylation of ATG16L1 S278, a site whose phosphorylation by ULK1 correlates with autophagic rate [48]. Importantly, to measure LC3b lipidation accurately, we pretreated cells with BafA1 to suppress the degradation of LC3b mediated by late-stage autophagic proteolysis. Note that since ATG16L1 S278 phosphorylation is not influenced by autophagic degradation [48], we analyzed ATG16L1 P-S278 preferentially in the absence of BafA1. To monitor AMPK activity, we measured the phosphorylation of ULK1 S555, a site phosphorylated directly by AMPK and proposed to induce autophagy [30,32]. In some experiments, we also monitored the phosphorylation of mTOR S1261, a site phosphorylated directly by AMPK [47]. To perform the amino acid withdrawal experiments, we refed cells with DMEM/FBS [10%] for 1 hr and then washed and refed the cells with amino acid-replete or amino acid-free DMEM containing glucose and dialyzed FBS (see Materials and Methods for more details). We therefore used identical media containing glucose and serum growth factors that contained or lacked only amino acids for the experiments in this study.

In wild type MEFs, HeLa, and C2C12 cells, amino acid withdrawal for 1-4 hr induced the accumulation of LC3b-II in the presence of BafA1, and amino acid withdrawal increased the phosphorylation of ATG16L1 S278 in both the absence and presence of BafA1, as expected (Fig. 1E-1G). The absence of AMPK ablated or reduced ULK1 S555 phosphorylation, thus demonstrating functional inactivation of AMPK (Fig. 1E-1G). Unexpectedly, however, the accumulation of LC3b-II was not impaired by AMPKα DKO/DKD in amino acid-deprived cells treated with BafA1 relative to control cells (Fig. 1E-1G). In addition, ATG16L1 P-S278 was higher (not lower) in MEFs and C2C12 cells lacking AMPK relative to control cells in the absence of BafA1 (Fig. 1E, 1G). These results indicate that autophagy remains intact in MEFs, HeLa, and C2C12 cells lacking AMPK. Moreover, the elevation of ATG16L1 P-S278 in MEFs and C2C12 cells (in the absence of BafA1) suggested that AMPKα DKO/DKD may increase the initiation of autophagy induced by amino acid withdrawal. As a control, we examined MEFs lacking ATG5, an essential autophagy protein important for the lipidation of LC3b. ATG5 knockout ablated the formation of lipidated LC3b-II, as expected, as well as the stability of ATG16L1 (Fig. S1A). We also used a different method to blunt autophagic flux, i.e., the lysosomal protease inhibitors E64D plus pepstatin A. Again, we found that the accumulation of LC3b-II was not impaired by the absence of AMPK in MEFs (Fig. S1B). These results indicate that AMPK is not required for autophagy induced by amino acid deprivation.

### The absence of AMPK increases autophagy induced by amino acid withdrawal

We next monitored autophagic responses quantitatively in MEFs and C2C12 cells containing or lacking AMPK. Consistent with the time course results, amino acid deprivation for 2 hr in the presence of BafA1 induced the accumulation of LC3b-II (normalized to α-tubulin) to a statistically similar extent in wild type vs. AMPKα DKO MEFs (Fig. 2A, 2B) and in control vs. AMPKα DKD C2C12 cells (Fig. 2C, 2D). We obtained similar results in CRISPR/Cas9-engineered MEFs, C2C12, HEK293, and U2OS cells lacking AMPKα1/α2 (Fig. S2A-S2D), indicating that AMPK is not required for autophagy induced by amino acid deprivation in several cell types. Finding that results obtained from freshly generated AMPKα DKO MEFs (Fig. S2A) phenocopy those obtained from established AMPKα DKO MEFs generated over a decade ago from genetically modified mice (Fig. 1E, 2A) [49] reassured us that the phenotype of intact autophagy in AMPKα DKO MEFs results from the lack of AMPK and not from cellular rewiring resulting from chronic knockout. Unexpectedly, we observed that amino acid withdrawal reduced (rather than increased) phosphorylation of ULK1 S555, a site phosphorylated by AMPK and proposed to initiate autophagy [30,32] in all five cell lines (i.e., MEFs; C2C12; HeLa; HEK293; U2OS) (Fig. 1, 2, S2).

**Figure 2.**
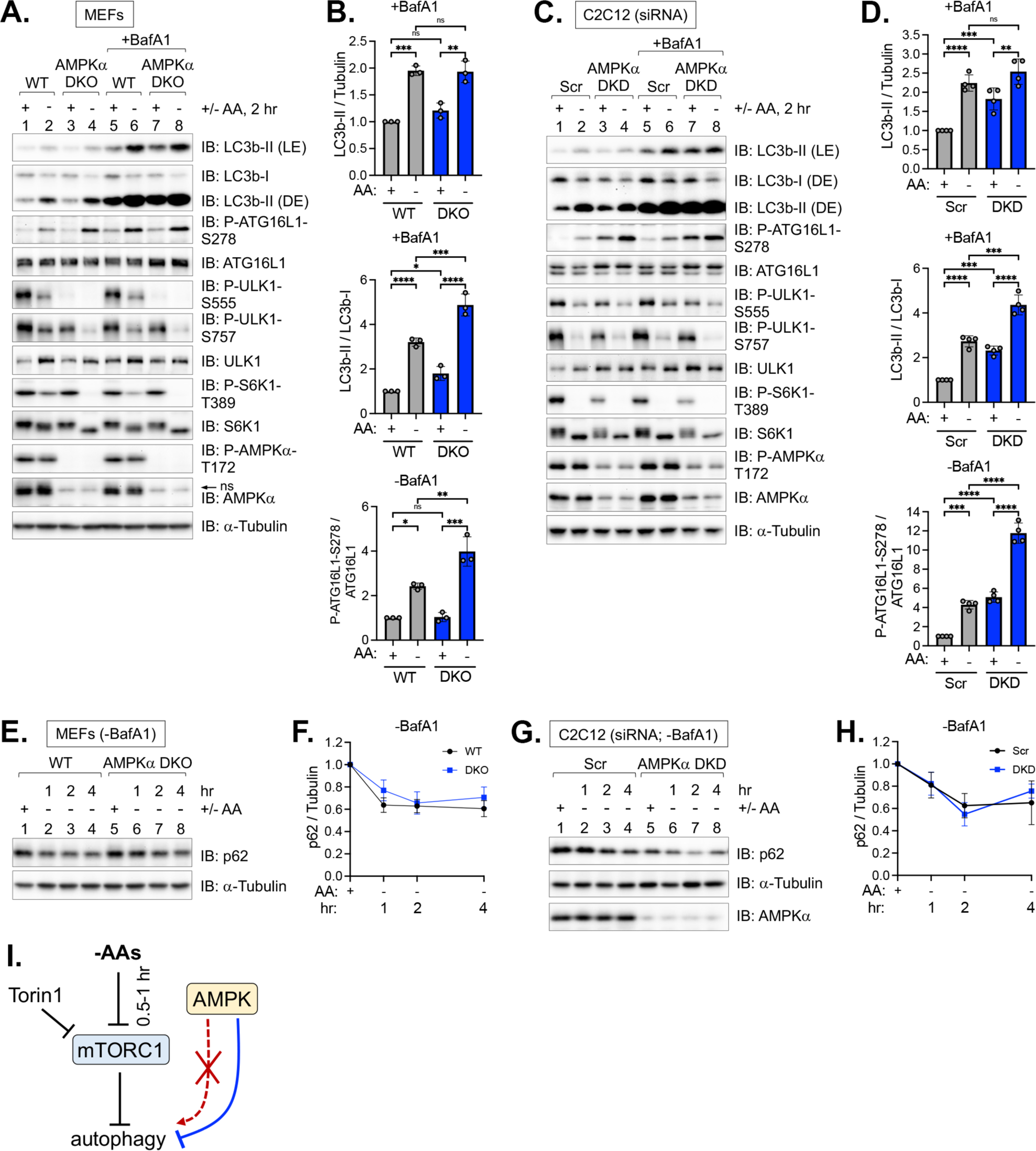
AMPK-deficient cells display increased autophagy induced by amino acid withdrawal. **(A)** WT and AMPKα DKO MEFs were fed with complete (+) or amino acid-free media (−) for 2 hr in the absence or presence of BafA1. ns, non-specific band detected by the AMPKα antibody in MEFs. **(B)** Quantification of LC3b-II/α-Tubulin, LC3b-II/LC3b-I, and ATG16L1 P-S278/ATG16L1 from (A). n = 3 independent experiments. **(C)** C2C12 cells were transiently transfected with scrambled or AMPKα1/α2 siRNA and treated as in (A). **(D)** Quantification of LC3b-II/ α-Tubulin, LC3b-II/ LC3b-I, and ATG16L1 P-S278/ATG16L1 from (C). n= 4 independent experiments. **(E)** WT and AMPKα DKO MEFs fed with complete (+) or amino acid-free media (−) for 1-4 hr in the absence of BafA1. The rate of degradation of p62/SQSTM1 was monitored by immunoblotting for p62. **(F)** Quantification of p62/α-Tubulin from (E). Normalized levels were set at 1.0 for wild type and AMPKα DKO MEFs. n = 3 independent experiments. **(G)** C2C12 cells were transiently transfected with siRNA as in (C) and treated as in (E) without BafA1. **(H)** Quantification of p62/α-Tubulin from (G). Normalized levels were set at 1.0 for control and AMPKα DKD C2C12 cells. n = 3 independent experiments. **(I)** Model: AMPK suppresses (blue line) rather than promotes (red line) autophagy upon amino acid deprivation. In complete media, mTORC1 suppresses the initiation of autophagy. The rapid inhibition of mTORC1 by acute withdrawal of amino acids or treatment with Torin1 initiates autophagy. Results in Fig. 1 and 2 indicate that AMPK is not required for autophagy induced by amino acid deprivation. In fact, AMPK may suppress autophagy in this context. Bars on graphs represent mean +/− SD (standard deviation). Two-way ANOVA was performed followed by Sidak’s multiple comparison post hoc tests; \**p* < 0.05; \*\**p* < 0.01; \*\*\**p* < 0.001; \*\*\*\**p* < 0.0001; ns, not significant. In (F) and (H), values represent mean +/− SD.

Upon induction of autophagy and lipidation of LC3b, levels of non-lipidated LC3b-I (upper band) decrease with corresponding increases in lipidated LC3b-II (lower band). The ratio of LC3b-II/LC3b-I (in the presence of BafA1) thus discriminates better between the induction of autophagy vs. a block in late-stage autophagic flux [19]. We found that LC3b-II/LC3b-I ratios were significantly higher in both complete media and amino acid-free media in AMPKα DKO MEFs and AMPKα DKD C2C12 cells relative to control cells (Fig. 2B, 2D). When we examined ULK1 signaling to ATG16L1 in the absence of BafA1, the induction of ATG16L1 P-S278 by amino acid withdrawal was not only unimpaired in AMPK-deficient MEFs and C2C12 cells but was increased to a significant extent (Fig. 2A-2D). In the basal state in the absence of BafA1, AMPK loss increased ATG16L1 P-S278 significantly in C2C12 cells (Fig. 2C, 2D)(see also Fig. 7C,7D, 7J, 7J) but not MEFs (Fig. 2A, 2B)(see also Fig. 7A, 7B, 7G,7H). To test whether these results extend to other ULK1 substrates, we examined the phosphorylation of ATG14 and Beclin-1. Like ATG16L1 P-S278, the loss of AMPK had no negative impact on the phosphorylation of ATG14 S29 or Beclin-1 S30 induced by amino acid withdrawal in MEFs and C2C12 cells (Fig. S2E, S2F). In fact, the loss of AMPK increased ATG14 P-S29 in both the basal and amino acid-deprived states (Fig. S2E, S2F). To complement our experiments utilizing genetic AMPK inactivation, we used the AMPK inhibitor BAY-3827 [50], which has improved specificity for AMPK over compound C, well-known for off-target effects [44]. We found that treatment of MEFs with BAY-3827 had no negative impact on the accumulation of LC3b-II or the phosphorylation of ATG16L1 S278 during amino acid deprivation (Fig. S2G). These results indicate that AMPK is not required for ULK1 signaling or LC3b lipidation upon amino acid deprivation. Moreover, they suggest that AMPK exerts a suppressive effect on these autophagic events.

p62/SQSTM1, an autophagy receptor that links cargo to growing phagophore membranes, becomes degraded by late-stage autophagic proteolysis [51]. We found that the degradation of p62/SQSTM1 induced amino acid withdrawal for 1-4 hr (in the absence of BafA1) occurred unimpaired in AMPK-deficient MEFs and C2C12 cells relative to control cells (Fig. 2E-2H). We noted that AMPKα DKO MEFs displayed elevated basal expression of p62 relative to control MEFs (Fig. 2E; also see Fig. 3E). As Sahani et al. demonstrated that prolonged amino acid deprivation transcriptionally upregulates p62, we speculate that chronic AMPK loss in MEFs may augment this response [52]. In addition, AMPKα DKD C2C12 cells displayed reduced basal expression of p62/SQSTM1 relative to control cells (Fig. 2G); we speculate that this phenotype results from increased basal autophagic proteolysis of p62/SQSTM1 in C2C12 cells lacking AMPK. To quantitate the rate of autophagic flux, we measured the fold change of LC3b-II/α-tubulin +BafA1/−BafA1 (i.e., the rate of LC3b turnover/degradation). The absence of AMPK increased autophagic flux significantly upon amino acid withdrawal in both MEFs and C2C12 cells (Fig. S2H, S2I). Together these results support a model whereby AMPK suppresses rather than promotes autophagy induced by amino acid deprivation (Fig. 2I).

**Figure 3.**
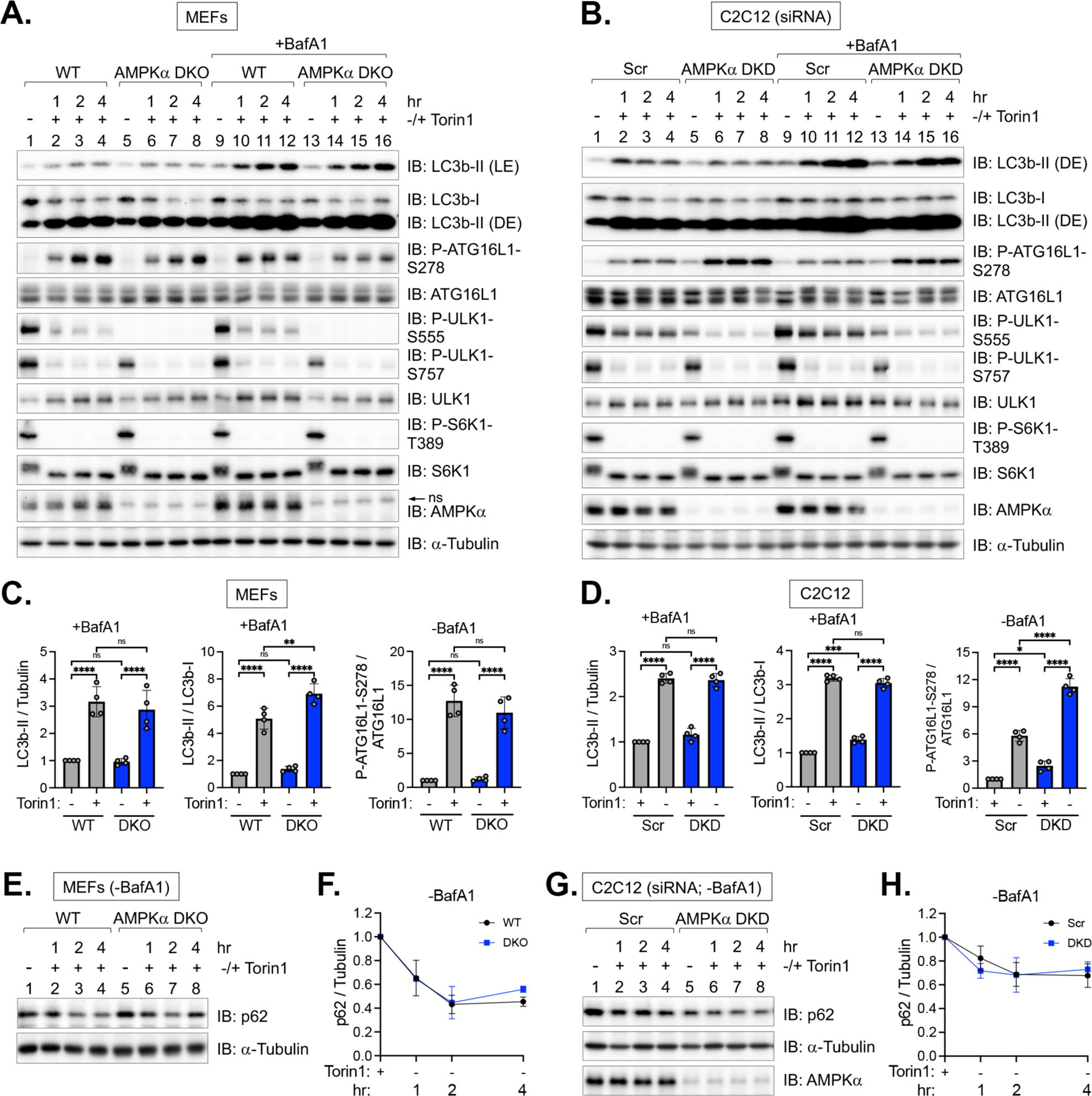
Autophagy induced by pharmacological mTORC1 inhibition occurs unimpaired in cells lacking AMPKα1/α2. **(A)** WT and AMPKα DKO MEFs were fed with complete media for 1 hr and then treated without (−) or with (+) Torin1 for 1-4 hr in the absence or presence of BafA1. Whole cell lysates were immunoblotted as indicated. LE: light exposure; DE: dark exposure; ns, non-specific band detected by the AMPKα antibody in MEFs. **(B)** C2C12 cells were transiently transfected with scrambled (Scr) or AMPKα1/α2 siRNA and treated as in (A). **(C)** Quantification of LC3b-II/α-Tubulin, LC3b-II/LC3b-I, and ATG16L1 P-S278/ATG16L1 from (A) (2 hr time point). n = 4 independent experiments. **(D)** Quantification of LC3b-II/α-Tubulin, LC3b-II/LC3b-I, and ATG16L1 P-S278/ATG16L1 from (B) (2 hr time point). n = 4 independent experiments. **(E)** WT and AMPKα DKO MEFs were treated as in (A) without BafA1, and levels of p62/SQSTM1 were measured by immunoblotting. **(F)** Quantitation of p62/α-Tubulin from (E). Normalized levels were set at 1.0 for wild type and AMPKα DKO MEFs. n = 3 independent experiments. **(G)** C2C12 cells were transiently transfected with siRNA as in (B), treated as in (A) without BafA1, and analyzed as in (E). **(H)** Quantitation of p62/α-Tubulin from (G). Normalized levels were set at 1.0 for control and AMPKα DKD C2C12 cells. n = 3 independent experiments. Bars on graphs represent mean +/− SD (standard deviation). Two-way ANOVA was performed followed by Sidak’s multiple comparison post hoc tests; \**p* < 0.05; \*\**p* < 0.01; \*\*\**p* < 0.001; \*\*\*\**p* < 0.0001; ns, not significant. In (F) and (H), values represent mean +/− SD.

### AMPK is not required for autophagy induced by pharmacological mTORC1 inhibition

Inhibition of mTORC1 by pharmacological agents induces autophagy to different extents depending on the type of inhibitor and cell type. For example, in most cell types, the active site mTOR inhibitor Torin1 induces autophagy robustly due to complete inhibition of mTORC1 while the allosteric inhibitor rapamycin induces autophagy weakly if at all [53,54]. To test whether AMPK is required for induction of autophagy in response to pharmacological mTORC1 inhibition, we treated MEFs and C2C12 cells containing or lacking AMPK with Torin1 for 1-4 hr and monitored autophagy in the absence and presence of BafA1. As expected, Torin1 reduced mTORC1 signaling strongly (as measured by S6K1 P-T389 and ULK1 P-S757) and induced a time-dependent accumulation of LC3b-II in BafA1-treated cells (Fig. 3A, 3B). Torin1 increased LC3b-II levels similarly in BafA1-treated MEFs (Fig. 3A, 3C) and C2C12 cells (Fig. 3B, 3D) containing and lacking AMPK, indicating that AMPK is not required for Torin1-induced autophagy. While Torin1 increased ATG16L1 P-S278 similarly in wild type and AMPKα DKO MEFs (no BafA1) (Fig. 3A, 3C), AMPKα DKD C2C12 cells displayed increased levels of ATG16L1 P-S278 relative to controls (no BafA1) (Fig. 3B, 3D), indicating elevated ULK1 signaling and increased initiation of autophagy. Interestingly, like amino acid withdrawal, Torin1 reduced the phosphorylation of ULK1 on the AMPK site (S555) in both MEFs and C2C12 cells (Fig. 3A, 3B). In addition, AMPK was not required for Torin1-induced autophagy in HeLa or HEK293 cells (Fig. S3A, S3B). We also confirmed that ATG5 KO MEFs display complete resistance to Torin1-induced autophagy (Fig. S3C). We next examined levels of p62/SQSTM1 in response to Torin1 treatment. p62/SQSTM1 degradation (in the absence of BafA1) was not impaired by loss of AMPK in MEFs (Fig. 3E, 3F) or C2C12 cells (Fig. 3G, 3H) relative to control cells in response to Torin1 treatment. Together these results demonstrate that Torin1-induced autophagy occurs unimpaired in the absence of AMPK, with evidence of increased autophagy.

### Formation of autophagosomal- and autolysosomal-like structures by amino acid withdrawal occurs unimpaired in cells lacking AMPK, with evidence of increased autophagy

Upon initiation of autophagy, lipidated LC3b, ATG16L1 P-S278, and p62/SQSTM1 concentrate on newly forming phagophore membranes that expand and mature into autophagosomes, which appear as puncta by confocal microscopy. In MEFs and C2C12 cells, AMPK loss had no negative impact on the formation of puncta containing these proteins in amino acid-deprived cells (Fig. 4A-4H). Moreover, in the absence of amino acids, AMPK loss increased the numbers of puncta containing LC3b significantly in C2C12 cells (but not in MEFs) (Fig. 4A-4D) while AMPK loss increased the numbers of puncta containing ATG16L1 P-S278 significantly in MEFs (but not C2C12 cells) (Fig. 4E-4H). In MEFs, AMPK loss increased the numbers of puncta containing p62/SQSTM1 significantly (Fig. 4I, 4J). The reason why AMPK loss differentially increases the numbers of LC3b and ATG16L1 P-S278 positive puncta depending on cell type remains unclear.

**Figure 4.**
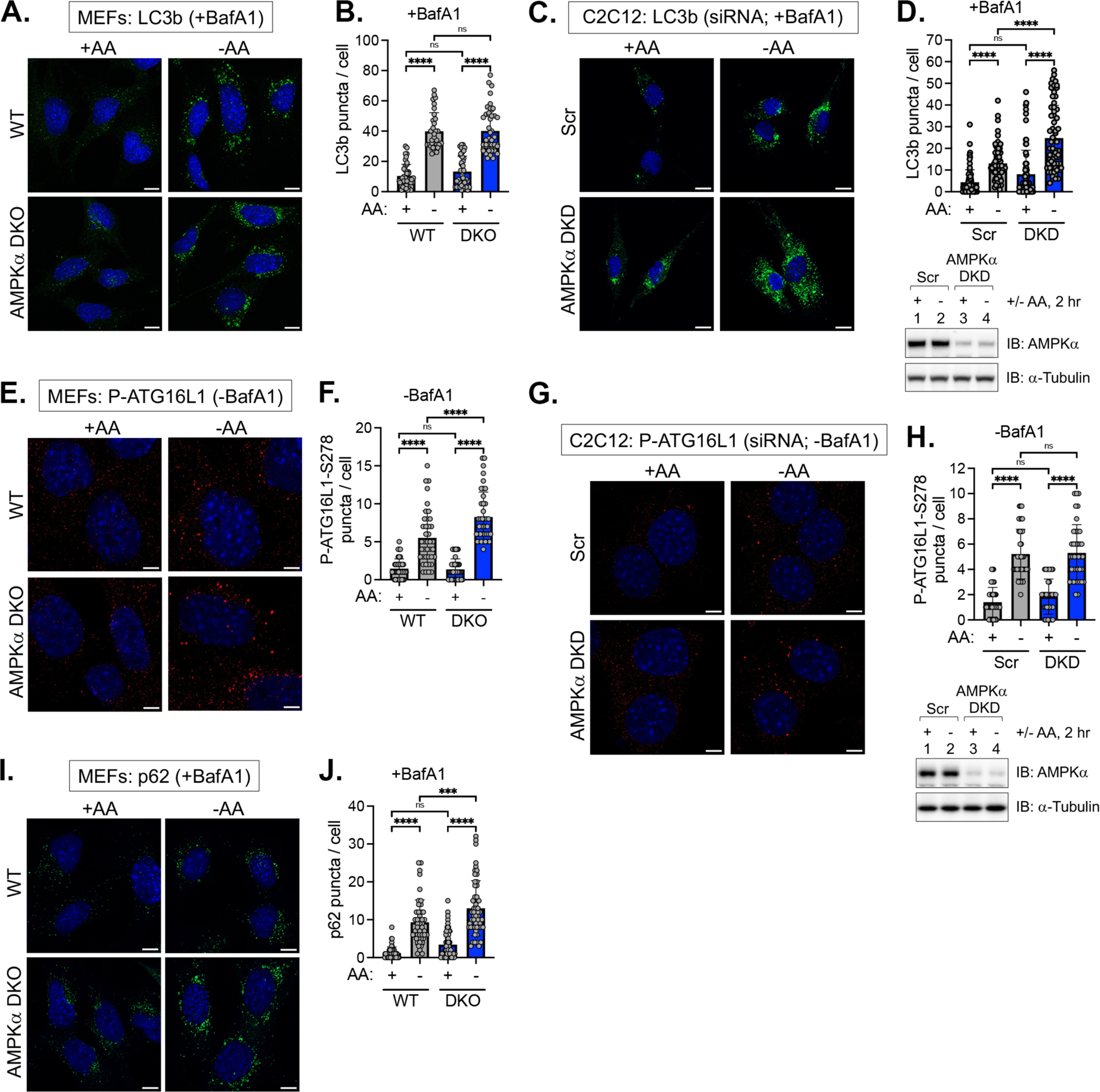
Induction of LC3b, ATG16L1 P-S278, and p62 positive puncta by amino acid deprivation occurs unimpaired in AMPK-deficient cells, with evidence of increased autophagy in the absence of AMPK. **(A)** WT and AMPKα DKO MEFs were fed with complete (+) or amino acid-free media (−) for 2 hr in the presence of BafA1. Cells were fixed and stained for endogenous LC3b. Scale bar: 10 μm. **(B)** Quantification of LC3b puncta number from (A). n = 30-40 cells from one experiment, representative of 4 independent experiments. **(C)** C2C12 cells were transiently transfected with scrambled (Scr) or AMPKα1/α2 siRNA and treated as in (A). Scale bar: 10 μm. **(D)** Quantification of LC3b puncta number from (C). n = 60-70 cells from two of four independent experiments. Below: Level of siRNA-mediated AMPKα knockdown in (C). **(E)** WT and AMPKα1/α2 DKO MEFs were fed with complete (+) or amino acid-free media (−) for 2 hr in the absence of BafA1. Cells were fixed and stained for endogenous ATG16L1 P-S278. Scale bar: 5 μm. **(F)** Quantification of ATG16L1 P-S278 puncta number from (E). n = 40 cells from two independent experiments. **(G)** C2C12 cells were transiently transfected with siRNA as in (C), fixed, and stained for endogenous ATG16L1 P-S278. Scale bar: 5 μm. **(H)** Quantification of ATG16L1 P-S278 puncta number from (G). n = 30-40 cells from one of two independent experiments. Below: Level of siRNA-mediated AMPKα knockdown in (G). **(I)** WT and AMPKα DKO MEFs were fed with complete (+) or amino acid-free media (−) for 2 hr in the presence of BafA1. Cells were fixed and stained for endogenous p62. Scale bar: 10 μm. **(J)** Quantification of p62 puncta from (I). n = 60-70 cells from 2 independent experiments. Bars on graphs represent mean +/− SD (standard deviation). Two-way ANOVA was performed followed by Sidak’s multiple comparison post hoc tests; \*\*\**p* < 0.001; \*\*\*\**p* < 0.0001; ns, not significant.

Transmission electron microscopy (TEM), the original method for monitoring autophagy, enables visual observation of autophagosome and autolysosome formation induced by amino acid withdrawal. We found that the ability of amino acid withdrawal to increase the number of autophagosomal- and autolysosomal-like structures was not impaired in MEFs or C2C12 cells lacking AMPK relative to control cells (Fig. 5A-5D). Moreover, the absence of AMPK increased the number of these structures upon amino acid withdrawal in MEFs (but not C2C12 cells) relative to control cells (Fig. 5A-5B). Together these results demonstrate that AMPK is not required for the formation of autophagosomal- and autolysosomal-like structures upon amino acid deprivation, with evidence of increased autophagy in the absence of AMPK.

**Figure 5.**
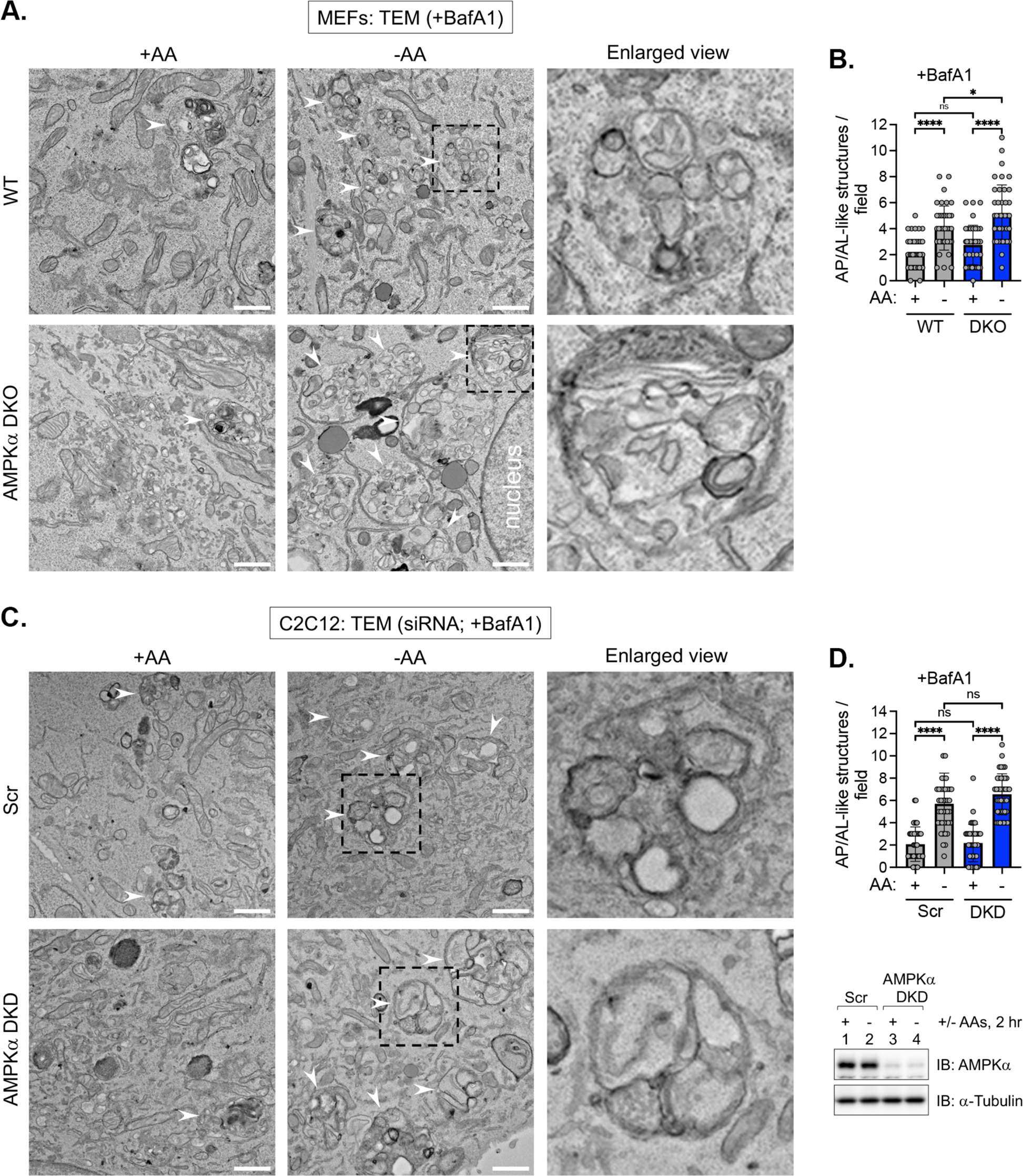
Formation of autophagosomal- and autolysosomal-like structures occurs unimpaired in AMPK-deficient cells upon amino acid deprivation. **A.** WT and AMPKα DKO MEFs were fed with complete (+ AA) or amino acid-free media (−AA) for 2 hr in the presence of BafA1. Cells were fixed, and the formation of autophagosomal- and autolysosomal-like structures was monitored by transmission electron microscopy (TEM). White arrowheads denote typical structures counted and quantified in B. An enlarged view of the dotted box from the middle panel (−AA) is shown in the panels on the right (Enlarged view). Scale bar: 1 μm. **B.** Quantification of the numbers of autophagosomal- and autolysosomal-like structures in (A). n = 40 fields of individual cells from one experiment. **C.** C2C12 cells were transiently transfected with scrambled (Scr) or AMPKα1/α2 siRNA and treated and analyzed as in (A). White arrowheads denote typical structures counted and quantified in D. An enlarged view of the dotted box from the middle panel (−AA) is shown in the panels on the right (Enlarged view). Scale bar: 1 μm. **D.** Quantification of the numbers of autophagosomal- and autolysosomal-like structures from (C). n = 40 fields of individual cells from one experiment. Right panel: Level of siRNA-mediated AMPKα knockdown in (C) was monitored by immunoblotting. Bars on graphs represent mean +/− SD (standard deviation). Two-way ANOVA was performed followed by Sidak’s multiple comparison post hoc tests; \**p* < 0.05; \*\*\*\**p* < 0.0001; ns, not significant.

### Pharmacological and physiological activation of AMPK suppresses autophagy induced by amino acid withdrawal

We next examined how the activation of AMPK affects basal and amino acid withdrawal-induced autophagy. To do so, we employed different agents to activate AMPK including the pharmacological agonist compound 991, glucose deprivation, and AICAR (an AMP mimetic compound). In MEFs treated with BafA1, compound 991 reduced the accumulation of LC3b-II (normalized to either α-tubulin or LC3b-I) induced by amino acid withdrawal in an AMPK-dependent and statistically significant manner (Fig. 6A, 6B). We obtained similar results in C2C12 cells in the amino acid-deprived state (Fig. 6C, 6D). Note that in C2C12 cells the accumulation of LC3b-II was AMPK-dependent with LC3b-II normalized to α-tubulin but not with LC3b-II normalized to LC3b-I, likely due to incomplete siRNA-mediated knockdown of AMPKα1/α2 (Fig 6D). In the basal state in MEFs, compound 991 treatment yielded variable effects on the accumulation of LC3b-II depending on the method of normalization; for example, 991 reduced the accumulation of LC3b-II significantly with LC3b-II normalized to α-tubulin but not with LC3b-II normalized to LC3b-I (Fig. 6A, 6B). In the basal state in C2C12 cells, however, compound 991 reduced LC3b significantly with LC3b normalized to either α-tubulin or LC3b-I (Fig. 6C, 6D). In HEK293 cells, 991 reduced the accumulation of LC3b-II in the basal and amino acid-deprived states in an AMPK-dependent manner (Fig. S4A). As expected, 991 increased ULK1 S555 phosphorylation in MEFs, C2C12 cells, and HEK293 cells, albeit modestly, indicating increased AMPK activity (Fig. 6A, 6C, S4A). To demonstrate unambiguously that 991 increased AMPK activity, we measured the phosphorylation of another AMPK substrate, i.e., mTOR S1261 [47]. Treatment with 991 increased mTOR P-S1261 in both the basal and amino acid-deprived states in an AMPK-dependent manner in MEFs and C2C12 cells (Fig. 6A, 6C). We also used AICAR to activate AMPK. AICAR suppressed both basal and amino acid withdrawal-induced accumulation of LC3b-II in an AMPK-dependent manner (Fig. S4B), which supports a prior study demonstrating that AICAR suppresses autophagy in rat hepatocytes [55]. We next examined the formation of LC3b puncta microscopically. Compound 991 reduced the number of LC3b puncta induced by amino acid withdrawal significantly in MEFs and C2C12 cells (Fig. 6E-6H). These results demonstrate that pharmacological AMPK activation with compound 991 suppresses autophagy induced by amino acid deprivation and likely also exerts a suppressive effect on basal autophagy.

**Figure 6.**
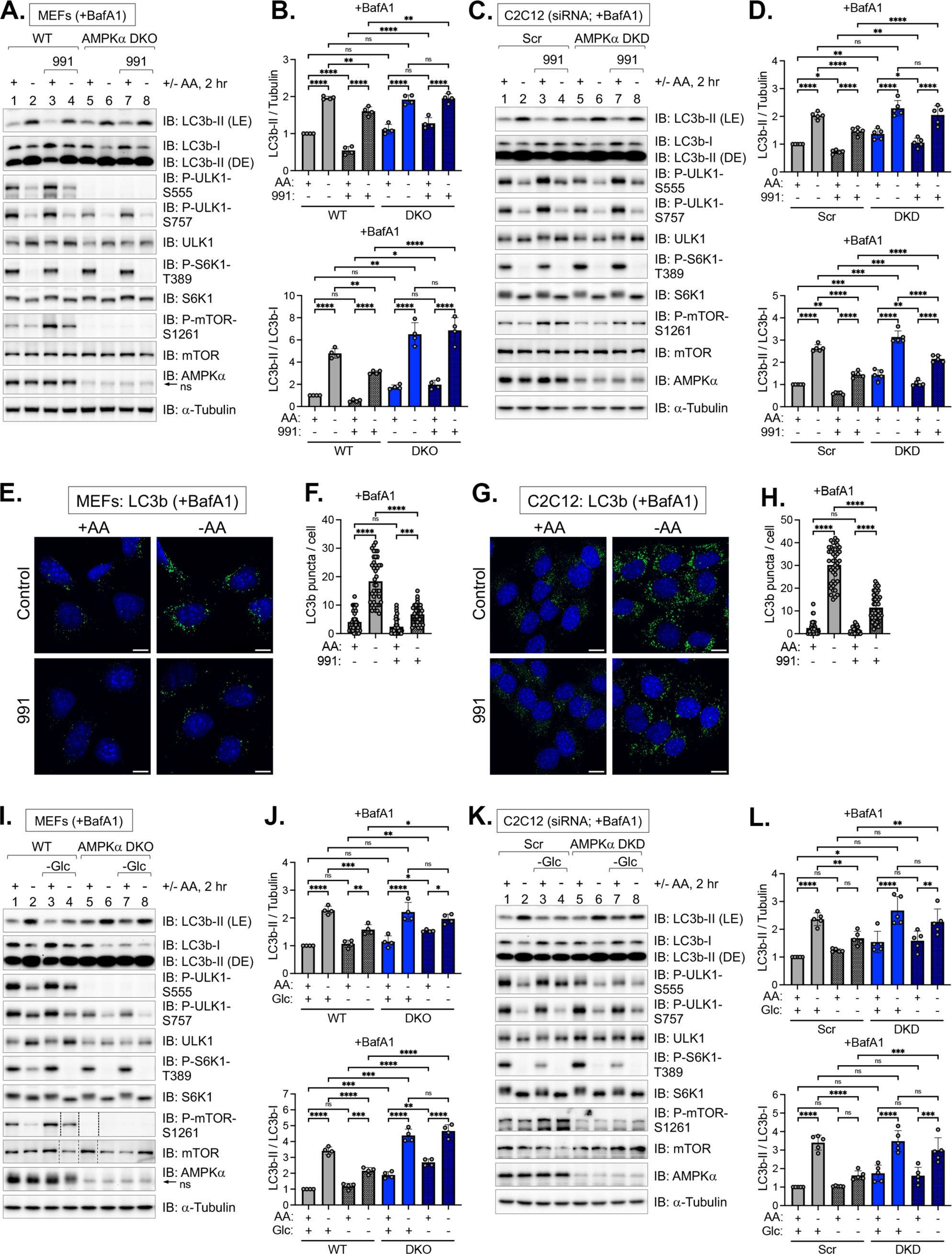
Activation of AMPK suppresses autophagy induced by amino acid deprivation. **(A)** WT and AMPKα DKO MEFs were pre-treated without or with compound 991 (30 min) and fed with complete (+) or amino acid-free media (−) for 2 hr in the absence or presence of 991 and the presence of BafA1. Whole cell lysates were immunoblotted as indicated. LE: light exposure; DE: dark exposure; ns, non-specific band detected by the AMPKα antibody in MEFs. **(B)** Quantification of LC3b-II/α-Tubulin (upper graph) and LC3b-II/LC3b-I (lower graph) from (A). n = 4 independent experiments. **(C)** C2C12 cells were transiently transfected with scrambled (Scr) or AMPKα1/α2 siRNA and treated as in (A). **(D)** Quantification of LC3b-II/α-Tubulin (upper graph) and LC3b-II/LC3b-I (lower graph) from (C). n = 5 independent experiments. **(E)** WT MEFs were pre-treated without or with 991 (30 min) and fed with complete (+) or amino acid-free (−) media in the absence or presence of 991 and presence of BafA1 for 2 hr. Cells were fixed and stained for endogenous LC3b. Scale bar: 10 μm. **(F)** Quantification of LC3b puncta number from (E). n = 40 cells from 2 independent experiments. **(G)** WT C2C12 cells were treated as in (E). Scale bar: 10 μm. **(H)** Quantification of LC3b puncta number from (G). n = 40 cells from 2 independent experiments. **(I)** WT and AMPKα DKO MEFs were fed with complete (+) or amino acid-free (−) media for 2 hr in the presence or absence of glucose (Glc) and the presence of BafA1. ns, non-specific band detected by the AMPKα antibody. Lanes #4 and #5 in the digital mTOR P-S1261 and mTOR images were re-ordered using Adobe Photoshop, as indicated by thin dotted lines. ns, non-specific band detected by the AMPKα antibody in MEFs. **(J)** Quantification of LC3b-II/α-Tubulin (upper graph) and LC3b-II/LC3b-I (lower graph) from (I). n = 4 independent experiments. **(K)** C2C12 cells were transiently transfected with siRNA as in (C) and treated as in (I). **(L)** Quantification of LC3b-II/α-Tubulin (upper graph) and LC3b-II/LC3b-I (lower graph l) from (K). n = 5 independent experiments. Bars on graphs represent mean +/− SD (standard deviation). Two-way ANOVA was performed followed by Sidak’s multiple comparison post hoc tests; \**p* < 0.05; \*\**p* < 0.01; \*\*\**p* < 0.001; \*\*\*\**p* < 0.0001; ns, not significant.

To increase AMPK activity more physiologically, we glucose-deprived cells. In agreement with results obtained using compound 991, withdrawal of glucose for 2 hr from BafA1-treated MEFs and C2C12 cells suppressed the ability of amino acid withdrawal to induce the accumulation of LC3b-II (normalized to α-tubulin or LC3b-I) to a significant extent in an AMPK-dependent manner (Fig. 6I-6L). Furthermore, glucose starvation alone (i.e., in the presence of amino acids) failed to increase the accumulation of LC3b-II in wild type MEFs or C2C12 cells (Fig. 6I-6L), indicating that glucose deprivation does not induce autophagy. This finding opposes the prevailing view that glucose deprivation induces autophagy. Curiously, the withdrawal of glucose from AMPKα DKO MEFs increased the accumulation of LC3b-II modestly but significantly when normalized to α-tubulin or LC3b-I (Fig. 6I, 6J). This observation agrees with Williams et al., who noted that shifting AMPKα DKO MEFs from high to low glucose induced the formation of LC3 puncta [56]. Thus, glucose deprivation can augment autophagy in the absence of the suppressive effect of AMPK in some cell types. Together these results demonstrate that physiological AMPK activation suppresses autophagy induced by amino acid deprivation. In addition, they demonstrate that glucose deprivation alone suppresses rather than induces autophagy.

### Activation of AMPK suppresses ULK1 signaling induced by amino acid withdrawal while AMPK loss increases lysosomal acidification

We next explored mechanisms by which AMPK suppresses autophagy. During amino acid sufficiency, mTORC1 phosphorylates ULK1 S757 (human S758), which suppresses the activity of the ULK1 complex [2,3,10,29]. Amino acid deprivation therefore initiates autophagy by relieving the suppressive effect of mTORC1 on ULK1. The activated ULK1 complex phosphorylates downstream substrates, including ATG16L1 (a component of the LC3b lipidation machinery) and ATG14 and Beclin-1 (components of the Vps34 complex) [2,3]. Amino acid deprivation also acidifies lysosomes independently of ULK1 [37], which facilitates autophagic degradation within autolysosomes. As the absence of AMPK increased ULK1 signaling to ATG16L1 during amino acid deprivation (see Fig. 2A-2D), we examined how activation of AMPK with compound 991 or glucose deprivation affects the activity of the ULK1 complex by measuring ATG16L1 P-S278, ATG14 P-S29, and Beclin-1 P-S30 in the absence of BafA1. In MEFs, compound 991 suppressed the ability of amino acid withdrawal to increase ATG16L1 P-S278 significantly in an AMPK-dependent manner (Fig. 7A, 7B). We obtained similar results in C2C12 cells (Fig. 7C, 7D), although the suppressive effect of 991 on ATG16L1 P-S278 was not wholly AMPK-dependent, likely due to incomplete siRNA-mediated knockdown of AMPKα1/α2. Compound 991 also suppressed the phosphorylation of the ULK1 substrates ATG14 S29 and Beclin-1 S30 in MEFs (Fig. S5A) and C2C12 cells (Fig. S5B) in both the basal and amino acid-deprived states in an AMPK-dependent manner. In addition, we confirmed that compound 991 increased AMPK activity in MEFs and C2C12 cells, as monitored by a modest increase in ULK1 P-S555 but a stronger increase in mTOR P-S1261 in the presence and absence of amino acids (Fig. 7A, 7C). Importantly, we confirmed the dependency of ATG16L1 P-S278 on ULK1 activity. Pretreating MEFs with the ULK1 inhibitor MRT-68921 (Fig. 7E, 7F) or knocking out ULK1 and its homologue ULK2 from MEFs (Fig. S5C) ablated ATG16L1 P-S278. These results confirm that the ATG16L1 P-S278 antibody indeed monitors ULK1 activity. Like compound 991, glucose withdrawal suppressed the ability of amino acid withdrawal to increase ATG16L1 P-S278 significantly in an AMPK-dependent manner and increased AMPK-mediated mTOR P-S1261 in MEFs (Fig. 7G, 7H) and C2C12 cells (Fig. 7I, 7J). These results demonstrate that pharmacological and physiological AMPK activation suppresses ULK1 signaling induced by amino acid deprivation. In addition, they demonstrate that glucose deprivation suppresses basal ULK1 signaling.

**Figure 7.**
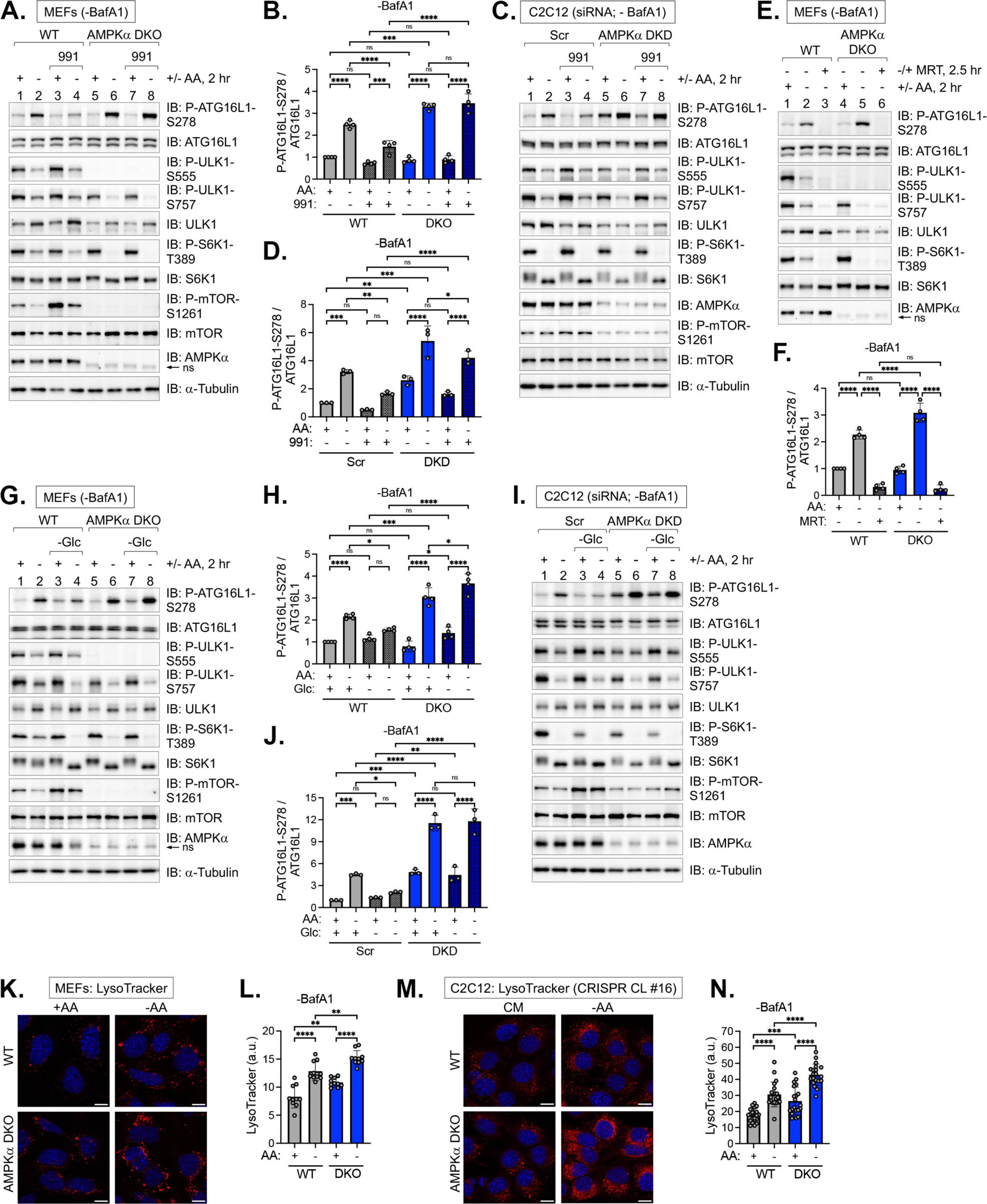
Activation of AMPK suppresses ULK1 signaling induced by amino acid deprivation while AMPK loss increases lysosomal acidification. **(A)** WT and AMPKα DKO MEFs were pre-treated without or with compound 991 (30 min) and refed with complete (+) or amino acid-free (−) media for 2 hr in the absence or presence of 991 and absence of BafA1. Whole cell lysates were immunoblotted as indicated. ns, non-specific band detected by the AMPKα antibody ns, non-specific band detected by the AMPKα antibody in MEFs. **(B)** Quantification of ATG16L1 P-S278/ATG16L1 from (A). n = 4 independent experiments. **(C)** C2C12 cells were transiently transfected with scrambled (Scr) or AMPKα1/α2 siRNA and treated as in (A). **(D)** Quantification of ATG16L1 P-S278/ATG16L1 from (C). n = 3 independent experiments. **(E)** WT and AMPKα DKO MEFs were pre-treated without or with MRT68921 (30 min) and re-fed with complete (+) or amino acid-free (−) media for 2 hr in the absence or presence of MRT68921 and absence of BafA1. **(F)** Quantification of ATG16L1 P-S278/ATG16L1 from (E). n = 4 independent experiments. **(G)** WT and AMPKα DKO MEFs were fed with complete (+) or amino acid-free (−) media in the presence or absence of glucose (Glc) for 2 hr and absence of BafA1. ns, non-specific band detected by the AMPKα antibody in MEFs. **(H)** Quantification of ATG16L1 P-S278/ATG16L1 from (G). n = 4 independent experiments. **(I)** C2C12 cells were transiently transfected with siRNA as in (C) and treated as in (G). **(J)** Quantification of ATG16L1 P-S278/ATG16L1 from (I). n = 3 independent experiments. **(K)** WT and AMPKα DKO MEFs were fed with complete (+) or amino acid-free (−) media for 2 hr in the absence of BafA1, treated with LysoTracker (30 min) and fixed. Scale bar: 10 μm. **(L)** Quantification of LysoTracker intensity from (K). n = 10 fields of cells with 4-6 cells/field (∼40-60 cells) from one experiment, representative of 2 independent experiments. **(M)** WT and AMPKα DKO C2C12 cells (CRISPR clone #16) were fed with complete (+) or amino acid-free (−) media for 2 hr in the absence of BafA1, treated with LysoTracker (30 min), and fixed. Scale bar: 10 μm. **(N)** Quantification of LysoTracker intensity from (M). n = 20 fields of cells with 15-20 cells/field (∼300-400 cells) from one experiment, representative of 2 independent experiments. Bars on graphs represent mean +/− SD (standard deviation). Two-way ANOVA was performed followed by Sidak’s multiple comparison post hoc tests; \**p* < 0.05; \*\**p* < 0.01; \*\*\**p* < 0.001; \*\*\*\**p* < 0.0001; ns, not significant.

As Nwadike et al. demonstrated that activation of AMPK with A-769662 suppresses the acidification of lysosomes mediated by amino acid withdrawal [37], we next examined lysosomal acidification in wild type and AMPK deficient cells by microscopic staining with LysoTracker, a cell-permeable, pH-sensitive dye that fluoresces more strongly in an acidic environment. We confirmed that amino acid withdrawal increases LysoTracker staining in MEFs (Fig. 7K, 7L) and C2C12 cells (Fig. 7M, 7N). Moreover, the absence of AMPK increased LysoTracker staining significantly in both complete and amino acid-free media (Fig. 7K-7N). These results confirm that AMPK suppresses lysosomal acidification in both the basal and amino acid-deprived states.

### Activation of AMPK suppresses ULK1 signaling and LC3b lipidation induced by pharmacological mTORC1 inhibition

We next tested how activation of AMPK with compound 991 affects autophagy induced by pharmacological mTORC1 inhibition compared to that induced by amino acid withdrawal. In complete media, Torin1 increased LC3b-II levels and ATG16L1 S278 phosphorylation to a level similar to or higher than that obtained in amino acid-free media in MEFs (Fig. 8A, 8B) and C2C12 cells (Fig. 8C, 8D). Treatment with 991 attenuated these Torin1-induced increases significantly (Fig. 8A-8D). Importantly, these Torin1-induced effects were AMPK-dependent, as 991 failed to reduce LC3b-II and ATG16L1 P-S278 significantly in AMPK-deficient MEFs and C2C12 cells cultured in complete media (Fig. 8E-8H). Although the allosteric, mTORC1-specific inhibitor rapamycin was less effective at inducing autophagy than Torin1 in complete media, particularly in C2C12 cells, compound 991 suppressed the rapamycin-induced increases in LC3b-II and ATG16L1 P-S278 (Fig. 8A-8D). These results indicate that activation of AMPK suppresses ULK1 kinase activity and LC3b lipidation induced by pharmacological mTORC1 inhibition. Interestingly, we also observed that Torin1 or rapamycin treatment attenuated the suppressive effect of 991 on basal and amino acid withdrawal-induced autophagy (Fig. 8A, 8C). These observations support a model whereby mTORC1 and AMPK each suppress autophagy in opposite directions through independent mechanisms operating in parallel (Fig. 8I).

**Figure 8.**
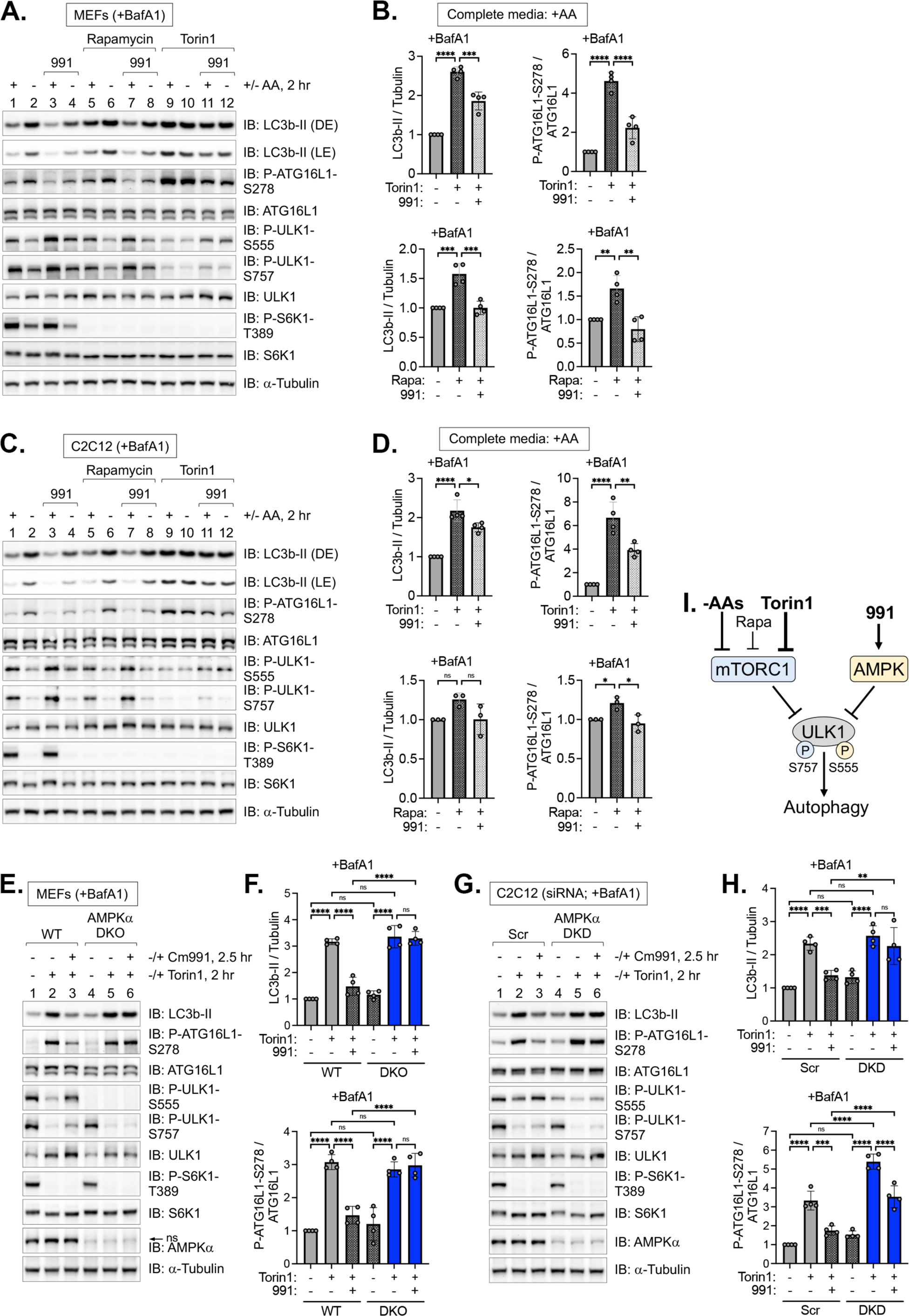
Activation of AMPK suppresses autophagic initiation and progression induced by pharmacological mTORC1 inhibition. **(A)** WT MEFs were pre-treated without or with rapamycin or Torin1 (30 min), then pre-treated without or with 991 (30 min), and then re-fed with complete (+) or amino acid-free (−) media lacking or containing the drugs above for 2 hr in the presence of BafA1. Whole cell lysates were immunoblotted as indicated (total treatment time, 3 hr). LE, light exposure; DE, dark exposure. **(B)** Quantification of LC3b-II/α-Tubulin (left panels) and ATG16L1 P-S278/ATG16L1 (right panels) from (A). Upper panels: Torin1-treated cells; lower panels: rapamycin-treated cells. n = 4 independent experiments. **(C)** WT C2C12 cells were treated as in (A). **(D)** Quantification of LC3b-II/α-Tubulin (left graphs) and ATG16L1 P-S278/ATG16L1 (right graphs) from (C). Upper panels: Torin1-treated cells; lower panels: rapamycin-treated cells. n = 4 independent experiments. **(E)** WT and AMPKα DKO MEFs were pre-treated without (−) or with (+) 991 (30 min) and then treated without (−) or with (+) Torin1 in the absence or presence of 991 for 2 hr and presence of BafA1. ns, non-specific band detected by the AMPKα antibody in MEFs. **(F)** Quantification of LC3b-II/α-tubulin and ATG16L1 P-S278/ATG16L1 from (E). n = 4 independent experiments. **(G)** C2C12 cells were transiently transfected with scrambled (Scr) or AMPKα1/α2 siRNA and treated as in (E). **(H)** Quantification of LC3b-II/α-tubulin and ATG16L1 P-S278/ATG16L1 from (G). n = 4 independent experiments. **(I)** Model: Activation of AMPK with compound 991 suppresses Torin1- and amino acid withdrawal-induced autophagy. Conversely, inactivation of mTORC1 with Torin1 attenuates the ability of AMPK to suppress autophagy. Bars on graphs represent mean +/− SD (standard deviation). One-way ANOVA was performed followed by Sidak’s multiple comparison post hoc tests; \**p* < 0.05; \*\**p* < 0.01; \*\*\**p* < 0.001; \*\*\*\**p* < 0.0001; ns, not significant.

### AMPK supports the reactivation of mTORC1 signaling induced by prolonged amino acid deprivation

Our results described above reveal that AMPK paradoxically suppresses rather than promotes autophagy during prolonged amino acid deprivation. We were therefore curious as to how AMPK controls mTORC1 signaling in this context, i.e., we asked how AMPK controls the reactivation of mTORC1 that occurs during prolonged amino acid deprivation (at ∼2-4 hr) after the initial inactivation of mTORC1 (at 0.5-1 hr). As AMPK inhibits mTORC1 during energetic stress [21,22], we presumed that cells lacking AMPK would display elevated mTORC1 signaling [11,14–16]. As expected, the acute removal of amino acids for 0.5 hr reduced mTORC1 signaling, as monitored by the mTORC1-mediated phosphorylation of S6K1 T389, and mTORC1 signaling reactivated partially at later time points, increasing slightly at 1 hr after amino acid withdrawal and increasing more strongly at 2 hr, which was maintained up to 4-6 hr (Fig. 9A). HeLa cells responded similarly but with slightly slower kinetics (Fig. 9B). Importantly, we confirmed that autophagic proteolysis and a rise in internal amino acids underlies the reactivation of mTORC1 signaling following prolonged amino acid deprivation, at least in part. Pretreating MEFs or HeLa cells with BafA1 led to the accumulation of lipidated LC3b-II in the basal and amino acid-deprived states and blunted the reactivation of mTORC1 (Fig. S6A, S6B). When we examined the reactivation of mTORC1 signaling in AMPK-deficient cells, however, we observed impaired reactivation of mTORC1 in AMPKα DKO MEFs and HeLa cells relative to control cells, opposite our expectations (Fig. 9A, 9B). This finding opposes the established view that AMPK inhibits mTORC1 universally in all contexts.

**Figure 9:**
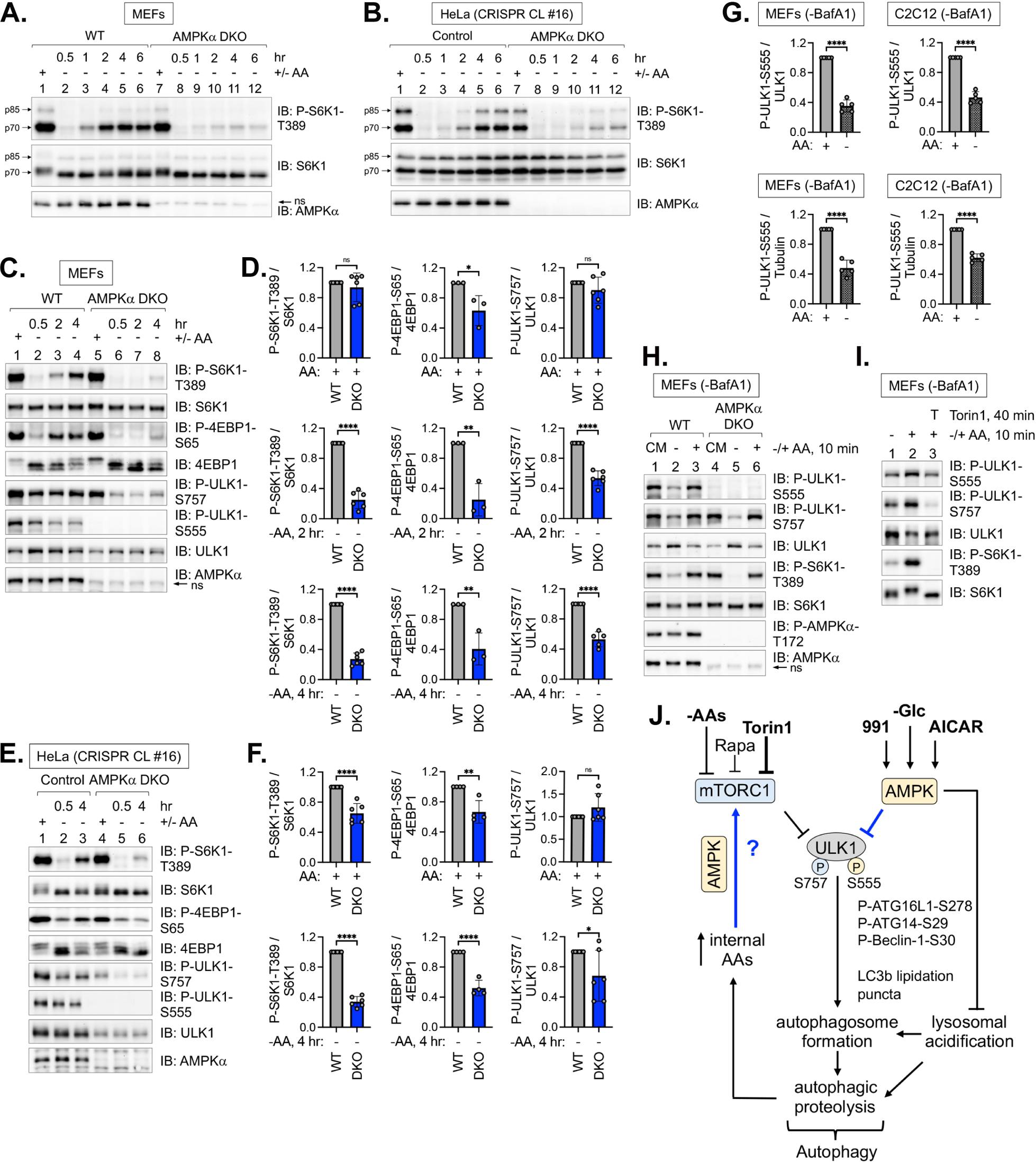
Impaired reactivation of mTORC1 signaling and reduced ULK1 S555 phosphorylation by the absence of AMPK during prolonged amino acid deprivation. **(A)** WT and AMPKα DKO MEFs were fed with complete media (+) for 1 hr or amino acid-free media (−) for 0.5-6 hr. Whole cell lysates were immunoblotted as indicated. ns, non-specific band detected by the AMPKα antibody. **(B)** Control and AMPKα DKO HeLa cells were treated as in (A). **(C)** WT and AMPKα DKO MEFs were fed with complete media (+) for 1 hr or amino acid-free media (−) for 0.5, 2, or 4 hr. Whole cell lysates were immunoblotted as indicated. ns, non-specific band detected by the AMPKα antibody in MEFs. **(D)** Quantification of S6K1 P-T389/S6K1 (left panels), 4EBP1 P-S65/4EBP1 (middle panels), and ULK1 P-S757/ULK1 (right panels) from (C). n = 3 or 6 independent experiments. **(E)** Control and AMPKα DKO HeLa cells (CRISPR clone #16) were treated as in (C). **(F)** S6K1 P-T389/S6K1 (left panels), 4EBP1 P-S65/4EBP1 (middle panels), and ULK1 P-S757/ULK1 (right panels) from (E). n = 4 or 6 independent experiments. **(G)** Quantification of ULK1 P-S555/total ULK1(upper panels) in MEFs or C2C12 cells cultured in complete media (+) or amino acid-free media (−) for 2 hr. n = 5 independent experiments. Quantification of ULK1 P-S555/α-tubulin (lower panels) in MEFs or C2C12 cells cultured in complete media (+) or amino acid-free media (−) for 2 hr. We normalized ULK P-S555/α-tubulin due to the extensive shifting that ULK1 undergoes on SDS-PAGE in response to amino acid levels (due to extensive phosphorylation), rendering it difficult to assess total ULK1 levels accurately. n = 5 independent experiments. **(H)** WT and AMPKα DKO MEFs were fed with complete media (CM) for 1 hr or amino acid-free media (−) for 50 min and stimulated without (−) or with (+) amino acids for 10 min. ns, non-specific band detected by the AMPKα antibody in MEFs. **(I)** WT and AMPKα DKO MEFs were amino acid-deprived (−) (50 min), pre-treated with Torin1 (30 min), and stimulated without (−) or with (+) amino acids (10 min). **(J)** Summary: Both mTORC1 and AMPK suppress autophagy. In amino acid-replete media, mTORC1 phosphorylates ULK1 S757 while AMPK phosphorylates ULK1 S555 (and likely other sites) to suppress the activity of the ULK1 complex. AMPK also suppresses the acidification of lysosomes. The inactivation of mTORC1 by amino acid withdrawal or Torin1 treatment reduces AMPK-mediated phosphorylation of ULK1. These events contribute to the activation of the ULK1 complex, which initiates the formation of autophagosomes. Upon induction of autophagy by amino acid deprivation or Torin1 treatment, the activation of AMPK by compound 991, glucose withdrawal, or AICAR suppresses autophagy. Upon prolonged amino acid withdrawal, autophagic proteolysis raises levels of internal amino acids, which reactivate mTORC1 in a manner partly dependent on AMPK. Bars on graphs represent mean +/− SD (standard deviation). Student’s t-tests (unpaired) were performed; \**p* < 0.05; \*\**p* < 0.01; \*\*\*\**p* < 0.0001; ns, not significant.

We next quantified the reactivation of mTORC1 signaling in cells containing or lacking AMPK. In MEFs, mTORC1-mediated signaling to S6K1 (P-T389) was significantly reduced by AMPKα DKO at 2 and 4 hr of amino acid withdrawal relative to wild type MEFs (Fig. 9C, 9D) (see also Fig. 1E, 2A, 2C, 7A, 7C, 7G; no BafA1 treatment). Similarly, AMPKα DKO HeLa cells displayed reduced reactivation of mTORC1 signaling toward S6K1 at 4 hr of amino acid withdrawal (Fig. 9E, 9F). We also asked whether AMPK supports mTORC1 signaling to 4EBP1, a well-characterized mTORC1 substrate that suppresses cap-dependent translation in the absence of mTORC1-mediated phosphorylation. In both MEFs and HeLa cells, mTORC1 signaling to 4EBP1 reactivated at 2-4 hr of amino acid withdrawal, as monitored by the phosphorylation of 4EBP1 S65 and by phosphorylation-mediated band-shifting (Fig. 9C-9F). In AMPKα DKO MEFs and HeLa cells, mTORC1-mediated phosphorylation of 4EBP1 S65 was impaired relative to control cells (Fig. 9C-9F). Thus, AMPK supports the reactivation of mTORC1 signaling toward two well-characterized substrates, i.e., S6K1 and 4EBP1, during prolonged amino acid deprivation. We also noted throughout this study that the absence of AMPK reduced the phosphorylation of ULK1 S757, another mTORC1 substrate, in amino acid-deprived cells (see Fig 1, 2, 7, and 9; no BafA1 treatment). The absence of AMPK also reduced total levels of ULK1, however (see Fig 1, 2, 7, and 9). When we normalized ULK1 P-S757 to total ULK1, we found that AMPK loss indeed reduced mTORC1 signaling to ULK1 significantly in both MEFs and HeLa cells, like mTORC1 signaling to S6K1 and 4EBP1 (Fig. 9C-9F). For reasons that remain unclear, mTORC1 signaling reactivates more strongly to S6K1 and 4EBP1 than to ULK1 during prolonged amino acid deprivation, which likely reflects important physiology. In addition, mTORC1 signaling to S6K1 reactivates minimally if at all in C2C12 cells. Together these results indicate that AMPK supports mTORC1 signaling toward at least three substrates, i.e., S6K1, 4EBP1, and ULK1, during prolonged amino acid deprivation (Fig. 9C-9F), revealing a previously unknown, non-canonical role for AMPK in the regulation of mTORC1.

### mTORC1 promotes AMPK-mediated ULK1 S555 phosphorylation

Throughout this study, we noted paradoxically that amino acid withdrawal (see Fig. 2, 6-9, S2) or Torin1 treatment (see Fig. 3 and 8) reduced (rather than increased) the phosphorylation of ULK1 S555, a site phosphorylated by AMPK and proposed to initiate autophagy. Earlier studies made a similar observation [37,39]. When we quantified the effect of amino acid levels on ULK1 S555 phosphorylation in MEFs and C2C12 cells, amino acid withdrawal reduced ULK1 P-S555 significantly (Fig. 9G). We also normalized ULK P-S555 to α-tubulin rather than to total ULK1 because ULK1 downshifts on SDS-PAGE in the absence of amino acids (due to reduced phosphorylation), thus generating a darker ULK1 band that when quantified would erroneously increase the fold reduction of ULK1 P-S555 caused by amino acid withdrawal. After normalizing ULK P-S555 to α-tubulin, amino acid withdrawal still decreased ULK P-S555 to a significant degree (Fig. 9G). As expected, AMPKα DKO or DKD ablated or reduced ULK1 P-S555 in MEFs and C212 cells, respectively (see Fig. 1-3), while AMPK activation with compound 991 increased ULK1 P-S555, albeit modestly (see Fig. 6-9). Since amino acid withdrawal or Torin1 treatment reduced ULK1 P-S555, we were curious as to how acute amino acid stimulation after acute amino acid withdrawal would affect ULK1 P-S555. The acute re-addition of amino acids (for 10 minutes) to MEFs deprived of amino acids (for 50 minutes) increased ULK1 P-S555 in an AMPK-dependent (Fig. 9H) and Torin1-sensitive (Fig. 9I) manner. As expected, amino acid stimulation increased S6K1 P-T389 and ULK1 P-S757. These results indicate that mTORC1 activity positively controls the ability of AMPK to phosphorylate ULK1 S555.

Together our results demonstrate that AMPK is not required for the induction of autophagy by amino acid deprivation or pharmacological mTORC1 inhibition. In fact, AMPK suppresses autophagy induced by amino acid deprivation and may also exert a modest suppressive effect on basal autophagy (Fig. 9J). The activation of AMPK by various methods (compound 991; glucose withdrawal; AICAR) suppressed autophagy induced by amino acid withdrawal or Torin1 treatment by reducing the activity of the ULK1 complex, as monitored by reduced phosphorylation of the ULK1 substrates ATG16L1 S278, ATG14 S29, and Beclin-1 S30 (Fig. 9J). Moreover, AMPK suppressed autophagosome formation, as monitored by reduced levels of lipidated LC3b-II, reduced numbers of LC3b, ATG16L1 P-S278, and p62 positive puncta, and reduced numbers of autophagosomal- and autolysosomal-like structures. AMPK also blunted the acidification of lysosomes in the basal and amino acid-deprived states, which impairs the fusion of autophagosomes with lysosomes (Fig. 9J). Mechanistically, AMPK suppresses autophagy by at least two mechanisms, i.e., by reducing ULK1 complex activity and blunting lysosomal acidification. Despite the potentially higher levels of internal amino acids generated by autophagic proteolysis in cells lacking AMPK relative to wild type cells, which could theoretically increase mTORC1 activity, mTORC1 reactivation was impaired during prolonged amino acid deprivation. These results indicate that AMPK supports the reactivation of mTORC1 signaling not by inducing autophagy but by an unknown mechanism (Fig. 9J). We speculate that AMPK-mediated support of mTORC1 signaling during amino acid deprivation may represent a third mechanism by which AMPK limits autophagy.

## Discussion

AMPK functions canonically as a sensor of energetic stress [24,25,27,46] that inhibits mTORC1 [21,22] and activates mTORC2 [47], resulting in improved energy balance and cell survival, respectively. The paradigm that AMPK promotes autophagy and mitophagy during glucose deprivation [9,29,30,32,33] fits well with these canonical functions of AMPK. Consistently, we found that AMPK-deficient cells displayed increased apoptotic cell death during nutrient stress caused by amino acid withdrawal (see Fig. 1A, 1B), but unexpectedly autophagy remained unimpaired (see Fig. 2-5). More surprisingly, we found that activation of AMPK suppressed autophagy induced by amino acid withdrawal (see Fig. 6-8). Our results thus present a revised model that challenges the prevailing view that AMPK promotes autophagy universally (Fig. 10A), an idea that has gained popularity over the last decade [2,3,27] since the discovery that AMPK phosphorylates ULK1 and Beclin-1 directly [29–31]. As autophagy consumes considerable amounts of ATP [57,58], the suppression of autophagy by AMPK aligns well with the canonical role of AMPK in the suppression of ATP-consuming processes. AMPK-mediated suppression of autophagy would enable low levels of metabolites and energy reserves to support essential cellular processes.

**Figure 10:**
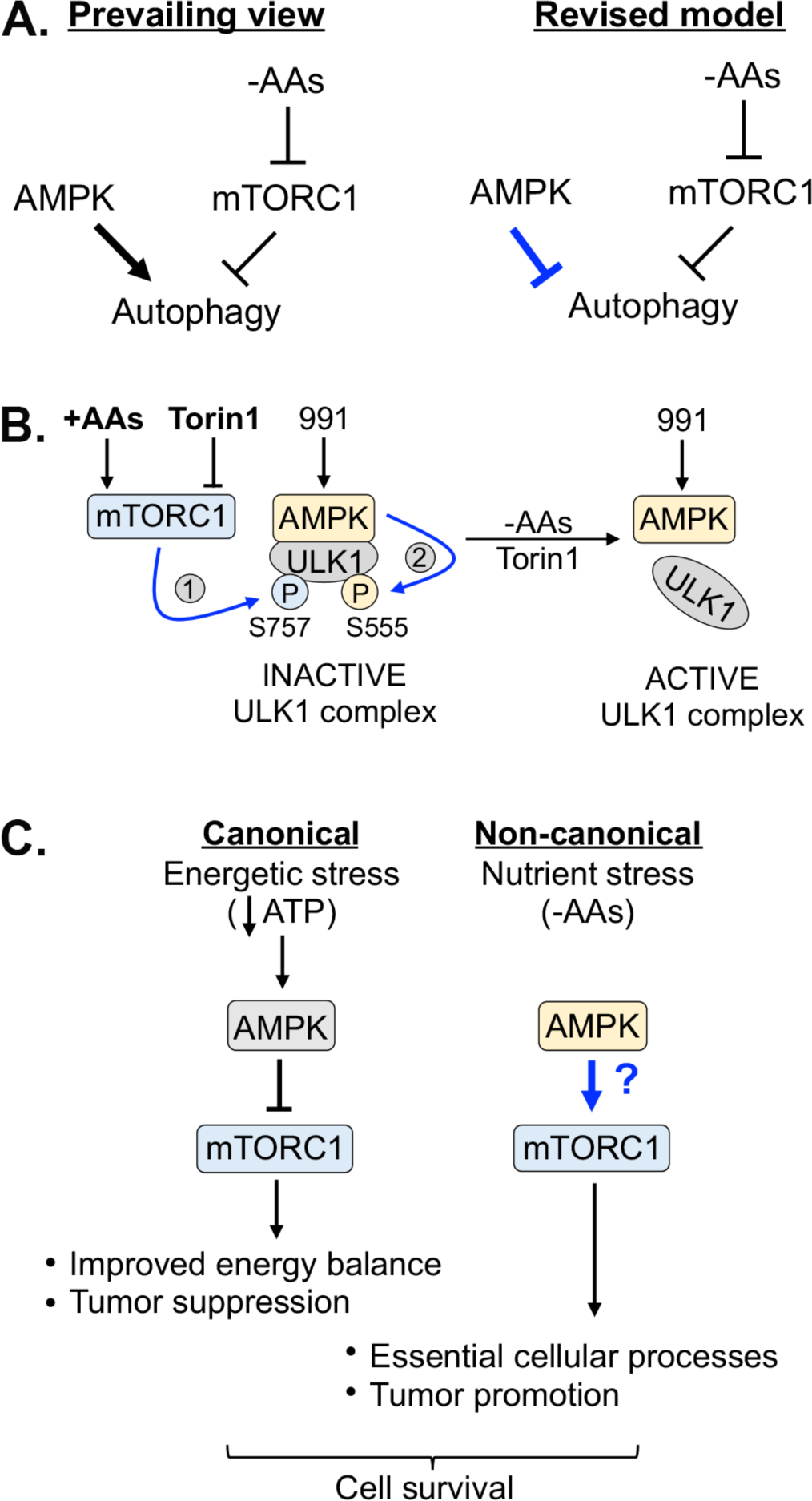
Models. **(A)** Prevailing view vs. revised model for how AMPK controls autophagy induced by amino acid withdrawal. **(B)** Model: mTORC1- and AMPK-mediated phosphorylation of ULK1 during amino acid sufficiency stabilizes the AMPK-ULK1 interaction. In complete media, active mTORC1 phosphorylates ULK1 S757, stabilizing the AMPK-ULK1 interaction and facilitating the AMPK-mediated phosphorylation of ULK1 S555. These events inactivate the ULK1 complex. Upon amino acid withdrawal or Torin1 treatment, the dephosphorylation of ULK1 S757 destabilizes the AMPK-ULK1 interaction, thus reducing the AMPK-mediated phosphorylation of ULK1 S555. These events activate the ULK1 complex. **(C)** Model: Depending on the cellular context, AMPK exerts different effects on mTORC1 signaling. During energetic stress, AMPK inhibits mTORC1 signaling and thereby suppresses anabolic metabolism, which improves energy balance to promote cell survival. During nutrient stress caused by amino acid deprivation, AMPK contributes to the reactivation of mTORC1 signaling, which supports essential cellular processes to promote cell survival.

While it makes physiological sense for glucose deprivation to induce autophagy and mitophagy to restore energy levels, we and other groups found that glucose withdrawal not only failed to induce the accumulation of LC3b-II in cells treated with a flux inhibitor [35–37] but suppressed the induction of autophagy upon amino acid withdrawal [37]. Furthermore, earlier studies found that activation of AMPK with AICAR or with the mitochondrial poisons phenformin or metformin either suppressed autophagy or had no effect [55,59]. Our results indicate that AICAR suppresses autophagy induced by amino acid withdrawal in an AMPK-dependent manner (see Fig. S4B). Williams et al. found that AMPKα DKO MEFs treated with a flux inhibitor displayed increased levels of LC3b-II in the basal and serum-deprived states relative to control cells [56]. Puente et al. showed that phosphorylation of ATG13 (a component of the ULK1 complex) on S258 and S224 by mTORC1 and AMPK, respectively, suppressed ULK1 signaling and the accumulation of LC3b-II during amino acid deprivation, suggesting that AMPK, like mTORC1, may suppress autophagy [60]. Therefore, several independent pieces of evidence suggest that AMPK activation or glucose deprivation fails to induce autophagy. As our study did not interrogate mitophagy, the role of AMPK in its control during amino acid or glucose deprivation warrants further investigation.

We and other groups found that amino acid withdrawal or pharmacological mTORC1 inactivation paradoxically reduced (rather than increased) the phosphorylation of ULK1 S555 [37,39,61], an AMPK site proposed to induce mitophagy upon glucose withdrawal [30,32]. Thus, mTORC1 positively controls ULK1 P-S555, but how? Shang et al. demonstrated that AMPK and ULK1 interact in complete media but dissociate upon the removal of amino acids [10,38] (Fig. 10B). Moreover, they found that blocking mTORC1-mediated phosphorylation of ULK1 S758 (mouse S757) using a non-phosphorylatable S758A mutant reduced the strength of the ULK1-AMPK interaction in complete media [10]. These results indicate that mTORC1-mediated phosphorylation of ULK1 S757 (S758) stabilizes the AMPK-ULK1 interaction (Fig. 10B). In addition, we found that the acute addition of amino acids to cells following acute amino acid withdrawal increased AMPK-mediated phosphorylation of ULK1 S555 in a Torin1 sensitive manner (see Fig. 9I). Together these results present a model whereby mTORC1-mediated phosphorylation of ULK1 S757 (S758) stabilizes the AMPK-ULK1 complex, which facilitates AMPK-mediated phosphorylation of ULK1 S555, possibly due to spatial proximity, to inhibit the ULK1 complex (Fig. 10B). We speculate that ULK1 S757 and S555 phosphorylation occur hierarchically, with AMPK-mediated ULK1 S555 phosphorylation requiring prior ULK1 S757 phosphorylation mediated by mTORC1. Why? Because inhibition of mTORC1 with amino acid withdrawal or Torin1 treatment reduced AMPK-mediated ULK1 P-S555 while Torin1 treatment blocked the ability of amino acid stimulation to increase AMPK-mediated ULK1 P-S555 (Fig. 9I; Fig. 10B).

Assays monitoring LC3b lipidation by western blot or the formation of LC3b and p62/SQSTM1 positive puncta by microscopy are prone to misinterpretation if not conducted in the presence of a flux inhibitor, which may explain why the concept that AMPK promotes autophagy has become an established view. For example, if the experimental design were to lack an autophagic flux inhibitor, and if one were to interpret the above autophagy readouts through the lens that AMPK promotes autophagy, then reduced levels of LC3b-II or LC3b- and p62-positive puncta upon inactivation of AMPK could be interpreted as “reduced” autophagy when in reality autophagy had been increased, leading to faster LC3b and p62 degradation by autolysosomes. We speculate that the absence of flux inhibitors may have led some groups to conclude that AMPK promotes autophagy. It is important to note that the analysis of autophagy in mice by using LC3b-II levels or LC3b- and/or p62-positive puncta as readouts becomes even more challenging due to the inability to easily inhibit autophagic flux *in vivo*.

While this manuscript was in preparation, a recent study reported findings overlapping with ours regarding the relationship between AMPK and autophagy [61]. By studying MEFs and HCT116 cells, Park et al. found that the activation of AMPK (using A769662, glucose withdrawal, or mitochondrial poisons) suppressed the ULK1-mediated phosphorylation of ATG14 S29 and Beclin-1 S30, the lipidation of LC3b, and the induction of LC3b and WIPI2 positive puncta induced by amino acid withdrawal [61]. Similarly, by studying MEFs and C2C12 cells, we found that the activation of AMPK (using compound 991, glucose withdrawal, or AICAR) suppressed ULK1-dependent phosphorylation of ATG16L1 S278, ATG14 S29, and Beclin-1 S30, the lipidation of LC3b, and the induction of LC3b and p62/SQSTM1 positive puncta. Thus, both groups agree that activation of AMPK suppresses autophagic initiation and progression during amino acid deprivation. Park et al. observed that amino acid withdrawal or Torin1 treatment reduced AMPK-mediated ULK1 S555 phosphorylation [61], like us and other groups [37,39]. Like Shang et al. [10], they demonstrated that amino acid withdrawal or Torin1 treatment destabilized the AMPK-ULK1 interaction due to reduced mTORC1-mediated phosphorylation of ULK1 S757. Park et al. also demonstrated that AMPK-mediated phosphorylation of ULK1 S556 (mouse S555) and T660 stabilizes the AMPK-ULK1 interaction and inhibits ULK1-mediated phosphorylation of ATG14 and Beclin-1, likely by suppressing the kinase activity of the ULK1 complex [61]. In agreement with Nwadike et al., we and Park et al. found that glucose deprivation not only fails to induce autophagy but suppresses autophagy induced by amino acid withdrawal [37,61]. We and Park et al. demonstrated further that glucose withdrawal suppresses autophagy induced by amino acid withdrawal in an AMPK-dependent manner [61]. In summary, we and Park et al. agree that the activation of AMPK suppresses rather than promotes the induction of autophagy induced by amino acid withdrawal [61].

Our results differ from the study by Park et al. in one respect, however. They found that the accumulation of LC3b-II upon amino acid withdrawal (2 hr) was impaired in AMPKα DKO MEFs relative to wild type MEFs treated with BafA1 [61]. In addition, they found that the lipid kinase activity of the Vps34 complex, as measured by an *in vitro* assay, was lower in AMPKα DKO MEFs relative to wild type MEFs. They, therefore, concluded that AMPK plays dual roles in the control of autophagy during amino acid deprivation, with the activation of AMPK suppressing autophagy and its presence supporting optimal autophagy. This latter idea aligns with the ability of AMPK to promote lysosomal biogenesis [62–65]. In contrast, we found that the accumulation of LC3b-II remained unimpaired in AMPKα DKO MEFs deprived of amino acids (2 hr) and treated with BafA1 (see Fig. 1E, 2A-B, 6A-B, S2I, S4B). In addition, we found that the accumulation of LC3b-II remained unimpaired in other AMPKα DKO cell lines (e.g., C2C12; HeLa; HEK293; U2OS) (see Fig. 1F, 1G, 2C-D, S2B, S2C, S2D). When we generated a new line of control vs. AMPKα DKO MEFs using CRISPR/Cas9, we again found that the accumulation of LC3b-II in the presence of BafA1 remained unimpaired in AMPKα DKO MEFs relative to control MEFs deprived of amino acids (see Fig. S2A). Moreover, when we normalized LC3b-II to LC3b-I, we found that AMPKα DKO MEFs and AMPKα DKD C2C12 cells converted LC3-I to LC3b-II faster than control cells upon amino acid withdrawal (see Fig. 2A-2D). The reason for the discrepancy between our results and those of Park et al. regarding the accumulation of LC3b-II in AMPKα DKO MEFs remains unclear. Interestingly, Park et al. found that AMPKα DKO MEFs and HCT116 cells displayed elevated ULK1-mediated phosphorylation of ATG14 S29 relative to control cells during amino acid withdrawal [61], similar to our data monitoring ATG16L1 P-S278, ATG14 P-S29, and Beclin-1 P-S30 in MEFs and C2C12 cells lacking AMPK (see Fig. 2C-2F). These latter results support the idea that AMPK loss upregulates autophagy in the amino acid-deprived state.

AMPK inhibits mTORC1 signaling in response to energetic stress to attenuate ATP-consuming cellular processes, thus balancing energy supply and demand [21,22,24,26] (Fig. 10C). Surprisingly, however, we found that during nutrient stress caused by prolonged amino acid deprivation, AMPK supports the reactivation of mTORC1 signaling, not by inducing autophagy but by a new albeit unknown mechanism (see Fig. 9A-9F; Fig. 10C). Thus, AMPK responds to cellular stress in both contexts but controls mTORC1 signaling in opposite directions (Fig. 10C). Our work therefore reveals an unexpected relationship between AMPK and mTORC1 that challenges the prevailing view that AMPK inhibits mTORC1 universally in all cellular contexts. We speculate that during amino acid deprivation, AMPK-mediated support of mTORC1 signaling maintains essential cellular processes necessary for cell survival (e.g., perhaps the translation of stress-induced genes) (Fig. 10C). Many important questions remain, however. How can AMPK inhibit mTORC1 during energetic stress yet support mTORC1 signaling during amino acid deprivation? Since AMPK can assemble into 12 potential isoforms due to its heterotrimeric nature and the diversity of its catalytic (α1-2) and regulatory subunits (β1-2; γ1-3), we hypothesize that distinct AMPK isoforms may control mTORC1 in different ways and in different contexts. Future work will be required to decipher the molecular mechanisms and AMPK isoforms underlying support of mTORC1 signaling in amino acid-deprived contexts. We speculate that AMPK-mediated support of mTORC1 signaling, in addition to AMPK-mediated support of mTORC2 signaling [47], may contribute to its paradoxical role as a tumor promoter [25,40–45].

In summary, our work reveals an unexpected role for AMPK in the suppression of autophagy induced by amino acid deprivation, which challenges the prevailing view that AMPK promotes autophagy universally. In addition, our work unveils a context-dependent, non-canonical role for AMPK in the support of mTORC1 signaling during nutrient stress caused by amino acid deprivation. These findings prompt a reevaluation of how AMPK and its control of autophagy and mTORC1 signaling maintain health and contribute to disease when dysregulated.

## Materials and Methods

### Materials

General chemicals were from Thermo Fisher Scientific or Millipore Sigma. NP40 and Brij35 detergents were from Pierce. Complete Protease Inhibitor Cocktail tablets (EDTA-free) (11836170001) and Immobilon-polyvinylidene difluoride (PVDF) membrane (0.45 μM) (88518) were from Thermo Scientific. PhosSTOP phosphatase inhibitor cocktail tablets were from Roche (04906845001). Reagents for enhanced chemiluminescence (ECL) were from either Alkali Scientific (Bright Star XR92), Advansta (Western Bright Sirius HRP substrate), or Bio-Rad (1705061). Every Blot Blocking Buffer (12010020) and 4x Laemmli Sample Buffer (1610747) were from Bio-Rad. Torin1 was a gift from Dr. David Sabatini [100 nM], Rapamycin was from Calbiochem (553210) [20 nM], compound 991 (ex229) was from Selleck (S8654) [10 μM], BAY-3827 was from Selleck (S9833) [1 μM], Bafilomycin A1 was from Cayman (11038) [1 μM], AICAR was from CST (9944) [2 mM], MRT68921 was from Tocris (57805) [10 μM], E64D was from Fisher (NC0132253) [10 μg/ml], and pepstatin A was from Sigma (11359053001) [10 μg/ml]. LysoTracker was from Invitrogen (L7528). ReadyProbe cell viability reagents were from Thermo Fisher Scientific (R37609).

### Antibodies

The following antibodies for western blots were from Cell Signaling Technology: LC3b (2775); AMPKα P-T172 (4188); pan AMPKα1/α2 (i.e., AMPKα) (2532); ULK1 P-S555 (5869); ULK1 P-S757 (23988); ULK1 (8054); ATG14 P-S29 (92340); ATG14 (96752); Beclin-1 P-S30 (35955); Beclin-1 (3495); p62/SQSTM1 (5114); S6K1 P-T389 (9234); cleaved caspase 3 (9664); cleaved PARP (9544); α-tubulin (2144); ATG5 (12994); and mTOR (2972). Antibodies for western blots also included the following: ATG16L1 P-S278 (Abcam, ab195242); ATG16L1 (Abcam, ab187671); S6K1 (amino acids 485-502; rat 70 kDa isoform) (custom-made, as described [66], and mTOR P-S1261 (custom-made, as described [66]. Donkey anti-rabbit-HRP (Jackson ImmunoResearch, 711-095-152) detected ECL-based western blot signals, we used. For confocal immunofluorescence microscopy, the following antibodies from Jackson ImmunoResearch were used: Alexa 488-conjugated anti-rabbit-HRP (711545152) and Alexa 594 conjugated anti-rat-HRP (112585167).

### Generation of AMPK**α**1/**α**2 DKO cells using CRISPR/Cas9-mediated genome editing

We used CRISPR/Cas9-mediated genome editing technology [67] to knockout AMPKα1 (*PRKAA1)* and AMPKα2 (*PRKAA2)* from HeLa cells, C2C12 cells, HEK293 cells, and MEFs. Four mouse-specific gRNAs in total were used for targeting *PRKAA1* (i.e., AMPKα1) and *PRKAA2* (i.e., AMPKα2) in MEFs and C2C12 cells, two gRNAs for AMPKα1 and AMPKα2 each. These sequences were as described [68]: *PRKAA1-* (A) GGCTGTCGCCATCTTTCTCC; (B) GAAGATCGGCCACTACATTC. *PRKAA2-* (A) TCAGCCATCTTCGGCGCGCG; (B) GAAGATCGGACACTACGTGC. Two human-specific gRNAs in total were used for targeting *PRKAA1* and *PRKAA2* in HeLa and HEK293 cells, one gRNA for AMPKα1 and one gRNA for AMPKα2. These sequences were as described [62]: *PRKAA1* - GTTATTGTCACAGGCATATGG; *PRKAA2-* GACAGGCATATGGTTGTCCAT. gRNAs were sub-cloned into vector pSpCas9(BB)-2A-Puro (PX459) (Addgene, 48139, deposited by the F. Zhang lab), and gRNA plasmids for targeting *PRKAA1* and *PRKAA2* were co-transfected into cells with TransIT-LT1 (Mirus, MIR 2305) according to the manufacturer’s instructions. PX459 lacking a guide RNA was used to generate “control” cells. Cells transfected with PX459 plasmids were selected with puromycin for 72 hr, and drug-resistant colonies (or pools of colonies, indicated in the figure) were isolated and screened for AMPKα1/α2 double knockout by western blotting. The following puromycin concentrations were used for drug selection of transfected CRISPR/Cas9 vectors: HeLa [2 μg/mL]; C2C12 [4 μg/mL]; HEK293 [2 μg/mL]; MEFs [8 μg/mL].

### Cell lines, cell culture, and cell treatments

Cell lines (MEFs; C2C12; HeLa; HEK293; U2OS) were cultured in DMEM containing high glucose [4.5 g/liter], glutamine [584 mg/liter] and sodium pyruvate [110 mg/liter] (Life Technologies/ Thermo Fisher, 11995-065) supplemented with 10% fetal bovine serum (FBS) (Life Technologies/ Thermo Fisher, 10347-028) and incubated at 37°C in a humidified atmosphere with 10% CO2. The wild type and AMPKα1/α2 DKO MEFs, originally isolated from transgenic mice by Dr. B. Viollet (Inserme, Paris, France) [49] were shared by Dr. R. Shaw (Salk Institute, San Diego, CA). The WT and AMPKα1/α2 DKO U2OS cells were generated and shared by Dr. Reuben Shaw [68]. The wild type and ATG5 KO MEFs were shared by Dr. N. Mizushima (Tokyo Medical University, Tokyo, Japan) [69].

To study the effects of amino acid deprivation, we first fed cells at ∼80% confluency with complete DMEM (containing FBS [10%], amino acids, and glucose) for 1 hr. We next washed cells with PBS containing Mg^2+^ and Ca^2+^ and refed them with precisely matched amino acid-replete DMEM (i.e., complete media) or amino-acid free DMEM containing dialyzed FBS [10%] and glucose [4.5 g/L] for various periods of time (0.5-16 hr) depending on the experiment. We prepared matched amino acid-replete and amino acid-free media in the following way: Amino acid-free DMEM (US Biologicals, D9800-13) containing 10% dialyzed FBS (Life Technologies, A33820-01) served as our amino acid deprivation media and as our base media for making complete media. To make complete media, we added a 50X RPMI 1640 amino acid solution (Sigma, R7131) supplemented with fresh glutamine ([300 mg/L] final) (Sigma, G8540) to the amino acid-free base media at 1x final amino acids. Importantly, before adding the 50x amino acid-glutamine solution to the amino acid-free DMEM, we adjusted the pH of this solution to 7.4, as the pH of the 50X RPMI amino acid solution is ∼10. Note that it was critical to adjust the pH of the amino acid solution, as the addition of an alkaline solution to cultured cells activates AMPK, mTORC1, and mTORC2 due to a rise in intracellular pH [70,71]. To suppress autophagic proteolysis, cells were treated without or with BafA1 [1 μM] for the period of amino acid withdrawal or Torin1 treatment (i.e., 1, 2, or 4 hr), depending on the experiment. To stimulate cells with amino acids, the RPMI amino acid solution with L-glutamine (adjusted to pH 7.4) was added to cells in amino acid-free media at a 1x final concentration (diluted from a 50X stock). For glucose deprivation experiments, DMEM containing dFBS [10%] but lacking amino acids and glucose (US Biologicals, D9800-28) was reconstituted with the amino acid [1x] solution containing L-glutamine, D-glucose [4.5 g/L] (Sigma, G8270), or both before each experiment.

Summary of final concentrations of reagents/drugs used in this study: Torin1 [100 nM]; compound 991 (ex229) [10 μM]; BAY-3827 [1 μM]; Bafilomycin A1 [1 μM]; AICAR [2 mM]; Rapamycin [20 nM]; MRT68921 [10 μM]; E64D [10 μg/ml]; pepstatin A [10 μg/ml].

### Cell lysis and immunoblotting

Cells were washed twice with ice-cold PBS, lysed in ice-cold buffer containing NP-40 [0.5%] and Brij35 [0.1%], and incubated on ice (15 min). Lysates were clarified by spinning at 16,100 x g for 5 min at 4°C, and the post-nuclear supernatants were collected. Bradford assay was used to normalize protein levels for immunoblot analyses. Samples were resolved on SDS-PAGE gels and transferred to PVDF membranes using the Bio-Rad Trans-Blot Turbo Transfer System according to the manufacturer’s instructions. Immunoblotting was performed by blocking PVDF membranes in Tris-buffered saline (TBS) pH 7.5 with 0.1% Tween-20 (TBST) containing 3% non-fat dry milk and incubating the membranes in TBST containing 2% bovine serum albumin (BSA) containing primary antibodies or secondary HRP-conjugated antibodies. ATG16L1 P-S278 antibody was diluted at 1:4000 in EveryBlot blocking buffer (Bio-Rad, 12010020) and incubated overnight. Western blots were developed by enhanced chemiluminescence (ECL) and detected digitally with a UVP ChemStudio Imaging System from Analytik Jena equipped with VisionWorks 9.2 software.

### RNA interference

AMPKα1 and α2 knockdown experiments in C2C12 cells were performed using scrambled control (Santa Cruz Biotechnology, sc-37007) or AMPKα1/α2-specific siRNAs (Santa Cruz Biotechnology, sc-45313) [10 nM]. C2C12 cells on 6-well plates were transiently transfected with RNAiMAX (Invitrogen, 13778-150) according to the manufacturer’s directions in two stages to prevent over-confluency at the time of cellular treatment and lysis. Cells were first transfected at ∼30-40% confluency and transfected again 24 hr later at 10-20% confluency. Cells were treated and lysed 72 hr after the first transfection at ∼80-95% confluency.

### Confocal immunofluorescence microscopy

Cells were seeded on glass coverslips in 6-well plates. Following various treatments, cells were washed with cold PBS^+^ and fixed with 3.7% formaldehyde for 10 min. Cells were washed twice with PBS^+^, permeabilized in 0.2% TX-100 for 5 min, washed twice with PBS^+^, and blocked in 0.2% fish skin gelatin (FSG) for 1 hr. Coverslips were inverted on primary antibodies (1:100) (rabbit anti-P-ATG16L1-S278, Abcam, ab195242; rabbit anti-p62, CST, 2321) diluted in PBS^+^ containing 0.2% FSG and incubated overnight. Cells were washed 3 times with PBS^+^, and coverslips were inverted on Alexa 594-conjugated anti-rabbit antibody (1:500) (Jackson, 711585152) or Alexa 488-conjugated anti-rabbit antibody (1:500) (Jackson, 711545152) diluted in PBS^+^/ 0.2% FSG, incubated for 1 hr, and washed 3 times with PBS^+^. Coverslips for LC3b were fixed with ice-cold methanol at −20°C for 15 min, washed 3 times with PBS^+^, and blocked in PBS^+^ with 0.2% FSG and 0.3% TX-100 for 1 hr. Coverslips were incubated overnight with 250 μL primary antibody (1:100) (Alexa 488-conjugated mouse anti-LC3b, CST, 98557) diluted in PBS^+^ containing 0.2% FSG and 0.3% TX-100 and washed 3 times with PBS^+^. LysoTracker experiments were performed by adding LysoTracker Red DND-99 (Invitrogen, L7528) (50 nM) to the media for the last 30 minutes of treatment. Coverslips were mounted on slides using Prolong Gold with DAPI (CST, 8961). Images were captured using a 40x objective lens on a Zeiss LSM800 Confocal Laser Scanning Microscope. ImageJ/ Fiji (National Institutes of Health) was used to quantitate the numbers of puncta/ cell for LC3b, ATG16L1 P-S278, and p62 staining and to quantitate LysoTracker staining intensity over a field of multiple cells.

### Transmission electron microscopy (TEM)

Cells were seeded on MatTek dishes. Following various treatments, cells were fixed overnight at 4°C in 2.5% glutaraldehyde in 0.1 M cacodylate buffer (pH 7.2) and washed with 0.1 M cacodylate buffer (pH 7.2) three times for 5 minutes each at room temperature. Cells were post-fixed in 1% K_4_Fe (CN)_6_ and 1% OsO_4_ in 0.1 M cacodylate buffer (pH 7.2) for 15 minutes at 4°C. Samples were washed three times for 3 minutes each with 0.1 M cacodylate buffer (pH 7.2) at room temperature, and then samples were washed with 0.1 M sodium acetate buffer (pH 5.2) three times for two minutes each at room temperature. Samples were en bloc stained with 2% uranyl acetate (in 0.1 M sodium acetate buffer (pH 5.2) for 15 minutes at room temperature. Samples were washed three times for 1 minute each with deionized water at room temperature. The cells were stained with Walton’s lead aspartate for 15 minutes at 60°C. Samples were washed three times for 3 minutes each with 0.1 M sodium acetate buffer (pH 5.2) at room temperature and three times with water for 3 minutes. The cells were dehydrated with a series of ethanol concentrations (10, 30, 50, 70, 80, 90, and 95%), 5 minutes each at 4°C, and finally, dehydrated twice with 100% ethanol for 5 minutes at room temperature. The cells were infiltrated with a 2:1 ratio of 100% ethanol to Araldite 502-Embed 812 for 30 minutes and 10 hr in a 1:2 ratio of 100% ethanol to Araldite 502-Embed 812 at room temperature. Finally, cells were infiltrated overnight with 100% Araldite 502/Embed 812. Cells were polymerized for 24 hr at 70°C and sectioned with a Leica UC7 ultramicrotome at 70 nm. The samples were imaged on a Jeol-1400 Plus Transmission Electron Microscope.

### Cell viability assay

Cells were seeded on 35 mm polylysine-coated glass bottom plates and amino acid-deprived for 16 hr. Cell viability was assessed by adding ReadyProbe reagents (Thermo Fisher Scientific, R37609) for the last 15 min of incubation according to the manufacturers’ directions. NucBlue Live reagent (Hoechst 33342) stains the nuclei of all cells, whereas NucGreen Dead stains the nuclei of cells with compromised plasma membranes (i.e., dead cells). Images were collected on an inverted epifluorescence microscope (Nikon TE2000E) equipped with a Photometrics CoolSnap HQ camera and analyzed using ImageJ/ Fiji (National Institutes of Health).

### Image editing

Adobe Photoshop was used to prepare digital western blot images for presentation, using only the levels, brightness, and contrast controllers equivalently over the entire image. In certain instances, lanes were reordered using Adobe Photoshop, as indicated by thin dashed lines in the western blot panels.

### Statistical analysis

Results are presented as mean ± standard deviation (SD). Data were analyzed by one-way or two-way ANOVA followed by Sidak’s multiple comparison post hoc tests or by unpaired Student’s t-tests, as indicated in the figure legends. Values of *p* <0.05 were considered significant. *\*p*<0.05; *\*\*p*<0.01; *\*\*\*p*<0.001; *\*\*\*\*p*<0.00001.

## Acknowledgments

We thank Dr. B. Viollet for the wild type and AMPKα DKO MEFs and Dr. R. Shaw for sharing these MEFs. We thank Dr. N. Mizushima for the wild type and ATG5 KO MEFs and Dr. JH Lee for sharing these MEFs. We thank Dr. D. Sabatini for sharing Torin1.

## Disclosure statement

We attest that we have no competing interests. Funders or other external entities played no role in the experimental design, execution, or analysis of this work.

## Funding

This work was supported by grants to DCF from the NIH (R01-GM137577) and by the Michigan Diabetes Research Center (MDRC) (NIH-P30-DK020572).

## Abbreviations

AAs: amino acids
ADP: adenosine diphosphate
AICAR: 5-aminoimidazole-4-carboxamide ribonucleotide
AMP: adenosine monophosphate
AMPK: AMP-activated protein kinase
ATG14: autophagy-related 14
ATG16L1: autophagy-related 16 like 1
ATG5: autophagy-related 5
BafA1: bafilomycin A1
DKD: double knockdown
DKO: double knockout
ECL: enhanced chemiluminescence
LC3b: microtubule-associated proteins 1A/1B light chain 3b
MEFs: mouse embryonic fibroblasts
mTORC1: mechanistic target of rapamycin complex 1
mTORC2: mechanistic target of rapamycin complex 2
p62: ubiquitin-binding protein p62, aka SQSTM1/Sequestosome 1
S6K1: ribosomal protein S6 kinase 1
4EBP1: eIF4E [eukaryotic initiation factor 4E] binding protein 1
TEM: transmission electron microscopy
ULK1: Unc-51-like kinase 1
Vps34: vacuolar protein sorting 34

## Supplemental Information

See Supplementary Materials for Figures S1-S6.

## Notes

### Competing Interest Statement

The authors have declared no competing interest.

### Summary of Updates

This version has been corrected for typos and unclear text. Figures 6, 7, and 9 as well as Supplementary Materials have also been updated to address reviewer comments.

