## Supplementary Material: Figures S1-S6 for "Unexpected roles for AMPK in the suppression of autophagy and the reactivation of mTORC1 signaling during prolonged amino acid deprivation"

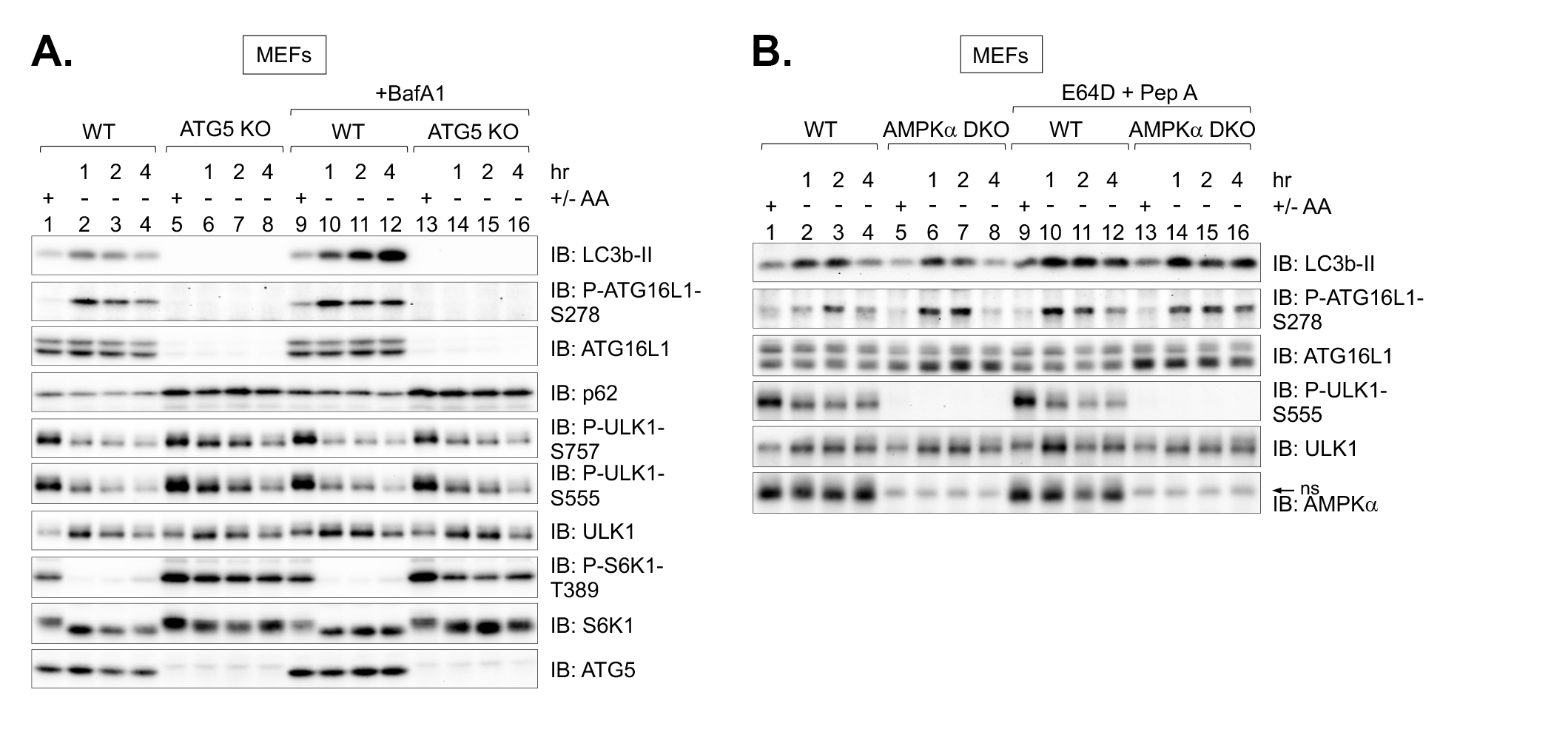

**Supplementary Figure S1 (related to Figure 1).** Autophagy induced by amino acid withdrawal requires ATG5 and occurs unimpaired in AMPKα DKO MEFs in the presence of protease inhibitors.

(**A**) WT and ATG5 KO MEFs were fed with complete media (+) for 1 hr or amino acid-free media (-) for 1-4 hr in the absence or presence of BafA1. Whole cell lysates were immunoblotted as indicated.

(**B**) WT and AMPKα DKO MEFs were treated as in (A) except that E64D and pepstatin A were used instead of BafA1.

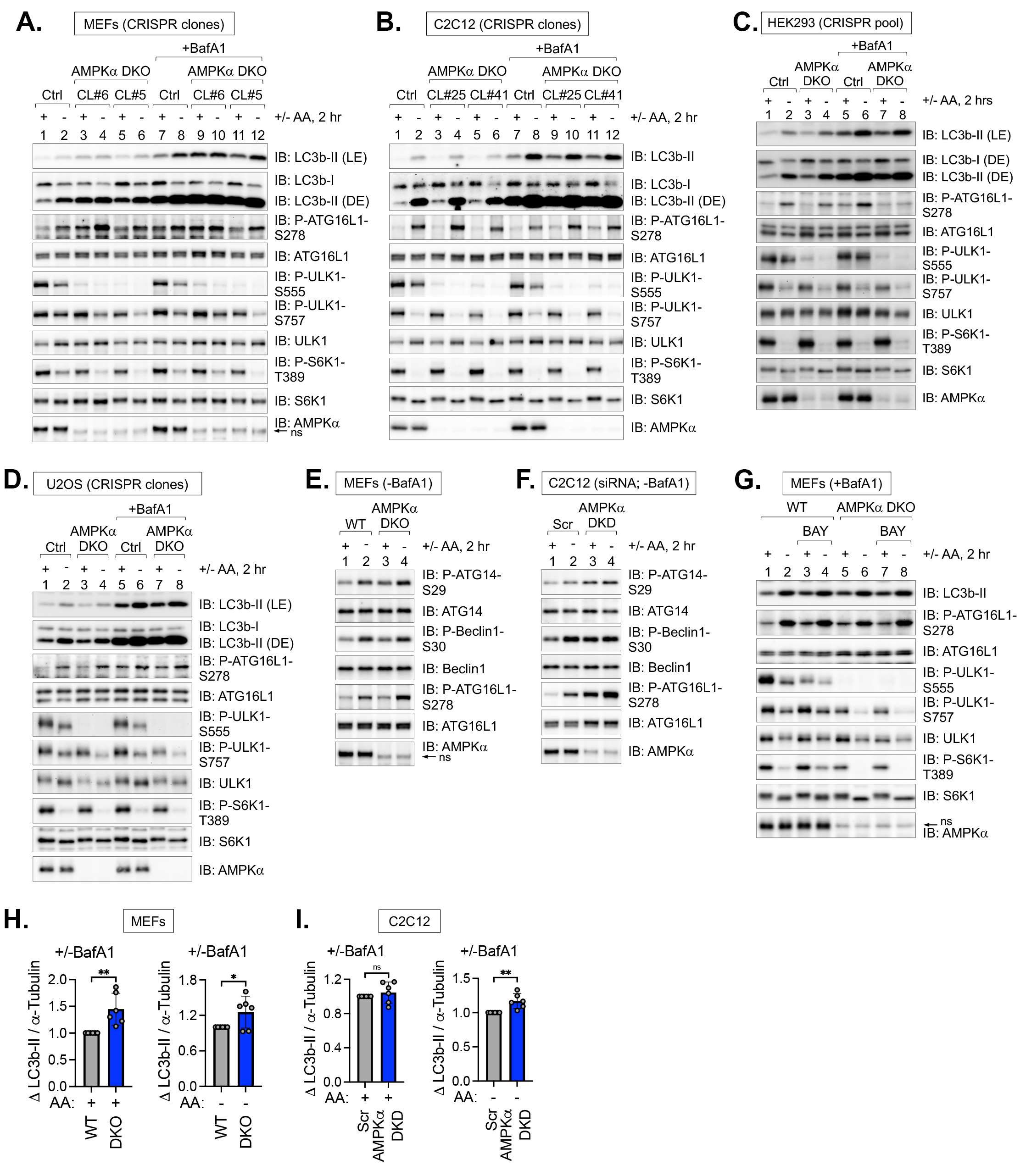

**Supplementary Figure S2 (related to Figure 2).** Autophagy induced by amino acid withdrawal occurs unimpaired in several cell lines lacking AMPKα1/α2

(**A**) Control (Ctrl) and AMPKα DKO MEFs (CRISPR clones #6 and #5) were fed with complete media (+) for 1 hr or amino acid-free (-) media for 2 hr in the absence or presence of BafA1. Whole cell lysates were immunoblotted as indicated. LE, light exposure; DE, dark exposure.

(**B**) Control (Ctrl) and AMPKα DKO C2C12 cells (CRISPR clones #25 and #41) were treated as in (A).

(**C**) Control (Ctrl) and AMPKα DKO HEK293 cells (CRISPR pool) were treated as in (A).

(**D**) Control (Ctrl) and AMPKα DKO U2OS cells (CRISPR pool) were treated as in (A).

(**E**) WT and AMPKα DKO MEFs were fed with complete (+) or amino acid-free media (-) for 2 hr in the absence of BafA1.

(**F**) C2C12 cells were transiently transfected with scrambled or AMPKα1/α2 siRNA and treated as in (E).

(**G**) WT and AMPKα DKO MEFs were pre-treated without or with BAY-3827 (BAY) (30 min) and fed with complete (+) or amino acid-free (-) media for 2 hr in the absence or presence of BAY-3827 and presence of BafA1.

(**H**) Quantification of LC3b-II (+BafA1)/ LC3b-II (-BafA1) normalized to α-Tubulin in MEFs in complete media (+) (left panel) and amino acid-free media (-) (right panel). n = 6 samples independent experiments.

(**I**) Quantification of LC3b-II (+BafA1)/ LC3b-II (-BafA1) normalized to α-Tubulin in C2C12 cells in complete media (+) (left panel) and amino acid-free media (-) (right panel). n = 6 independent experiments.

Bars on graphs represent mean +/- SD (standard deviation). Student’s t-test (unpaired) was performed; **p* < 0.05; ***p* < 0.01; ns: not significant.

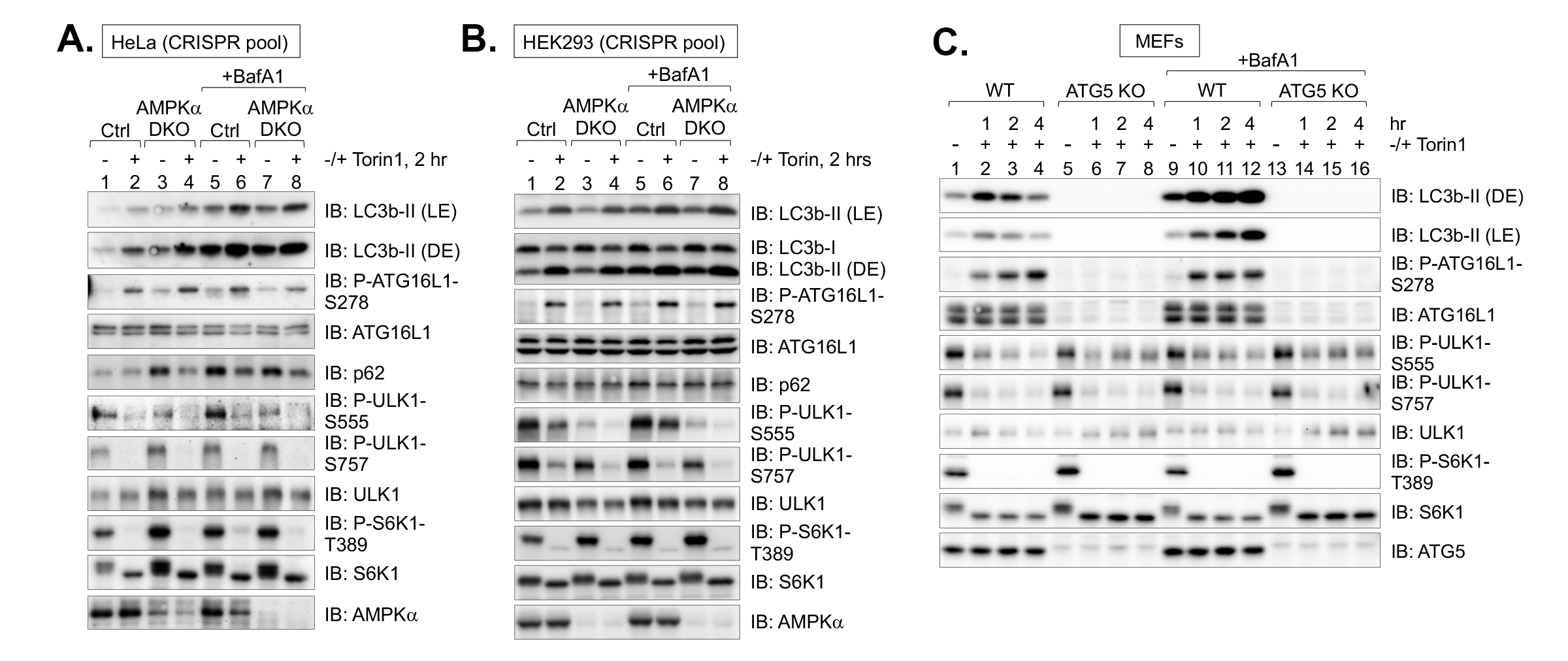

**Supplementary Figure S3 (related to Figure 3).** Autophagy induced by Torin1 occurs unimpaired in HeLa and HEK293 cells lacking AMPKα1/α2, and ATG5 KO abrogates Torin1-induced autophagy.

(**A**) Control (Ctrl) and AMPKα DKO HeLa cells (CRISPR pool) were treated without (-) or with (+) Torin1 for 2 hr in the absence or presence of BafA1. Whole cell lysates were immunoblotted as indicated. LE, light exposure; DE, dark exposure.

(**B**) Control (Ctrl) and AMPKα DKO HEK293 cells (CRISPR pool) were treated as in (A).

(**C**) WT and ATG5 KO MEFs were treated without (-) Torin1 for 1 hr or with (+) Torin1 for 1-4 hr in the absence or presence of BafA1.

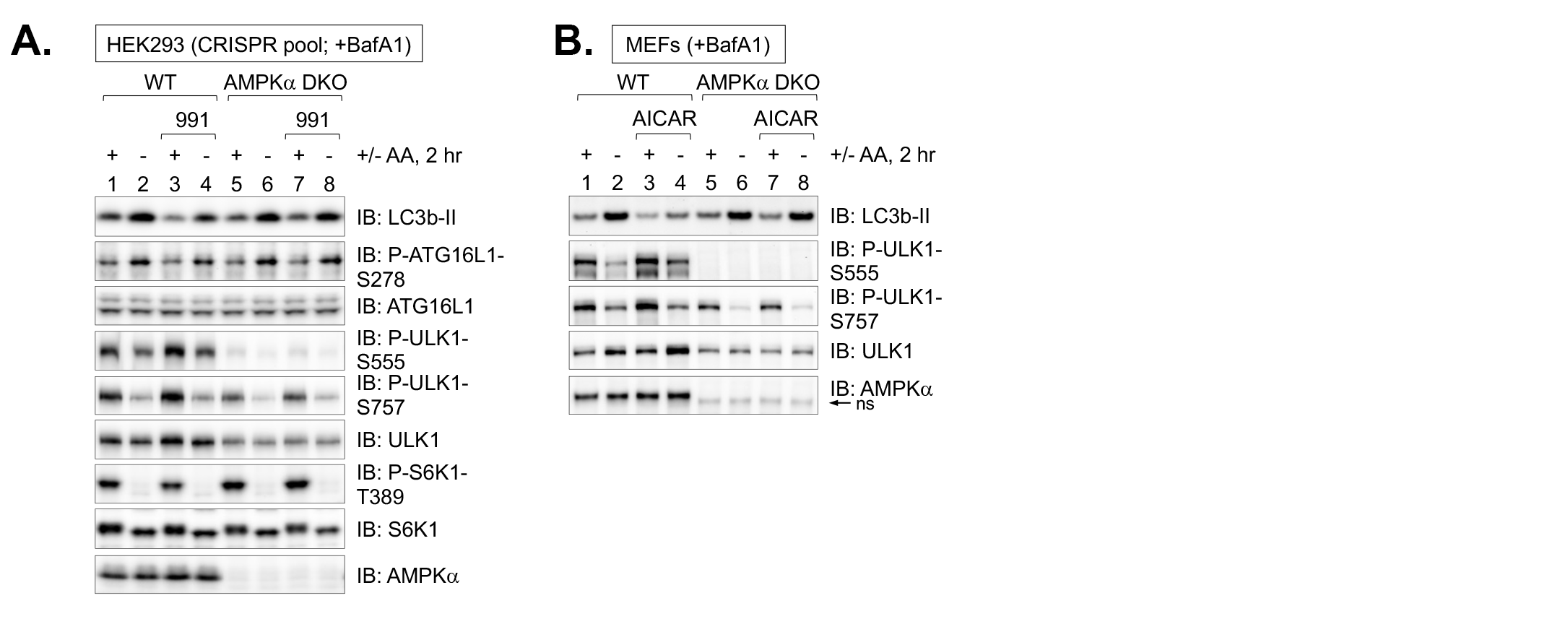

**Supplementary Figure S4 (related to Figure 6).** Activation of AMPK with 991 (in HEK293 cells) or AICAR (in MEFs) suppresses autophagy induced by amino acid withdrawal in an AMPK-dependent manner.

(**A**) Control (Ctrl) and AMPKα DKO HEK293 cells (CRISPR pool) were pre-treated without or with compound 991 (30 min) and fed with complete (+) or amino acid-free (-) media for 2 hr in the absence or presence of 991 and presence of BafA1. Whole cell lysates were immunoblotted as indicated.

(**B**) WT and AMPKα DKO MEFs were treated as in (C) except cells were treated with AICAR instead of 991.

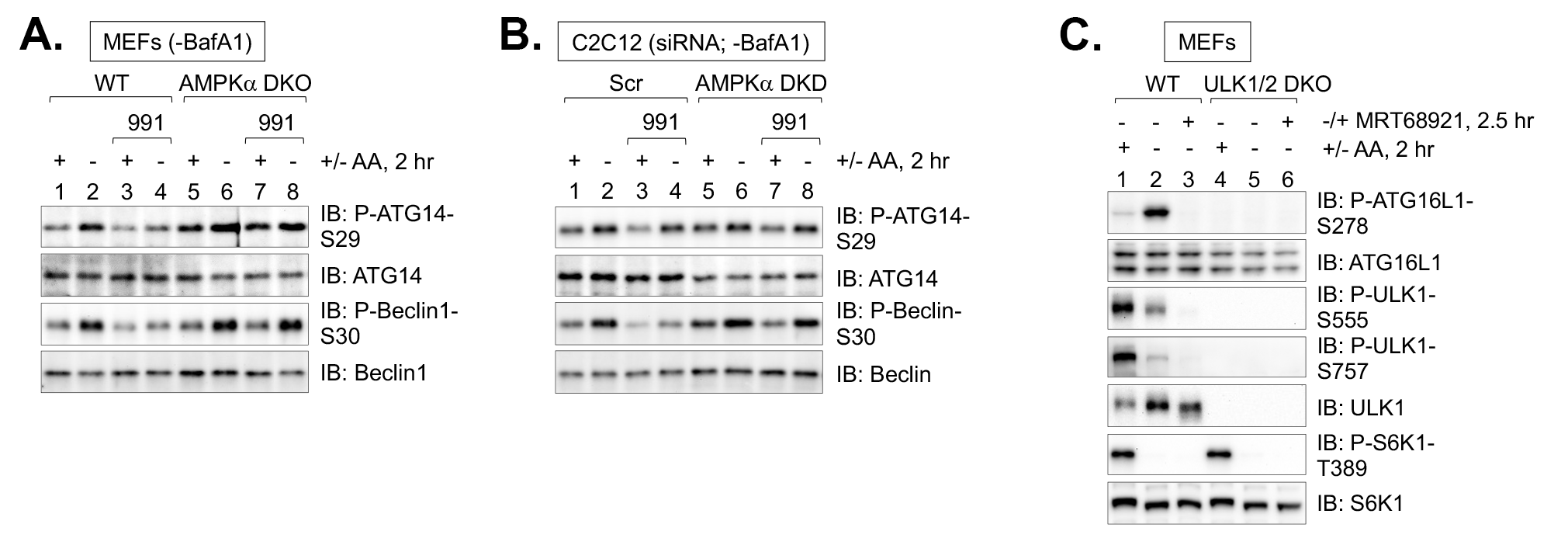

**Supplementary Figure S5 (related to Figure 7):** Activation of AMPK suppresses ULK1 activity towards other substrates (i.e., ATG14; Beclin1) while inactivation of ULK1/2 function inhibits autophagy initiation

(**A**) WT and AMPKα DKO MEFs were pre-treated without or with compound 991 (30 min) and refed with complete (+) or amino acid-free (-) media for 2 hr in the absence or presence of 991 and absence of BafA1 (same lysates as in Figure 7A). Whole cell lysates were immunoblotted as indicated.

(**B**) C2C12 cells were transiently transfected with scrambled (Scr) or AMPKα1/α2 siRNA and treated as in (A) (same lysates as in Figure 7C).

(**C**) WT and ULK1/2 DKO MEFs were pre-treated without or with MRT68921 (30 min) and fed with complete (+) or amino acid-free (-) media for 2 hr in the absence or presence of MRT68921 and absence of BafA1.

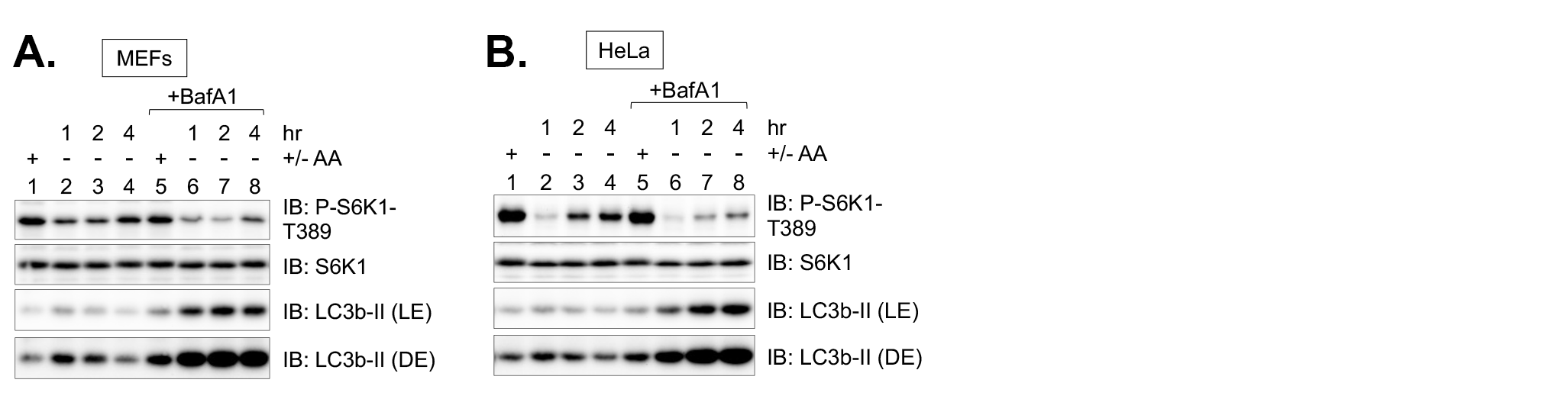

**Supplementary Figure S6 (related to Figure 9):** Inhibition of autophagic degradation blunts mTORC1 signaling and its reactivation during prolonged amino acid deprivation

(**A**) WT MEFs were fed with complete media (+) for 1 hr or amino acid-free media (-) for 1-4 hr in the absence or presence of BafA1. LE: light exposure; DE: dark exposure.

(**B**) WT HeLa cells were treated as in (A).
